# Circadian clocks in human cerebral organoids

**DOI:** 10.1101/2024.02.20.580978

**Authors:** Nina M Rzechorzek, Magdalena A Sutcliffe, Andrei Mihut, Koby Baranes, Nuzli Karam, Daniel Lloyd-Davies Sánchez, Sew Y Peak-Chew, Aiwei Zeng, Noah Poulin, Estere Seinkmane, Kaiser Karim, Christopher M Proctor, Mark Kotter, Madeline A Lancaster, Andrew D Beale

**Author notes:** Corresponding author, **Correspondence and requests for materials** should be addressed to Nina Rzechorzek. These authors contributed equally (Andrei Mihut, Koby Baranes, Mark Kotter, Madeline A Lancaster, Andrew D Beale). **Author note** This manuscript is currently undergoing peer review and the results may be updated in due course.

## Abstract

Circadian rhythms result from cell-intrinsic timing mechanisms that impact health and disease^1,2^. To date, however, neural circadian research has largely focused on the hypothalamic circuitry of nocturnal rodents^3^. Whether circadian rhythms exist in human brain cells is unknown. Here we show *bona fide* circadian rhythms in human neurons, glia, cerebral organoids, and cerebral organoid slices (ALI-COs)^4–8^. Human neural circadian rhythms are synchronised by physiological timing cues such as glucocorticoids and daily temperature cycles, and these rhythms are temperature-compensated across the range of normal human brain temperatures^9^. Astrocyte rhythms are phase-advanced relative to other cultures and they modulate neuronal clock responses to temperature shift. Cerebral organoid rhythms are more robust at physiological brain temperatures; the relative amplitude of these rhythms increases over time in culture and their resetting capacity recapitulates key neurodevelopmental transitions in glucocorticoid signalling^10–14^. Remarkably, organoid post-transcriptional bioluminescent clock reporter rhythms are retained even when those of their putative transcriptional drivers are indiscernible^15^, and electrophysiology recordings confirm circadian rhythms in functional activity of monocultures, organoids, and ALI-COs. Around one third of the cerebral organoid proteome and phosphoproteome are circadian-rhythmic, with temporal consolidation of disease-relevant neural processes. Finally, we show that human brain organoid rhythms can be modulated and disrupted by commonly used brain-permeant drugs and mistimed cortisol exposure, respectively. Our results demonstrate that human brain cells and tissues develop their own circadian oscillations and that canonical mechanisms of the circadian clockwork may be inadequate to explain these rhythmic phenomena. 2D and 3D human neural cultures represent complementary and tractable models for exploring the emergence, disruption, and mechanics of the circadian neural clockwork, with important implications for chronobiology, brain function, and brain health.

## Introduction

Physiology is cyclical, oscillating over timescales from milliseconds to months. Circadian oscillations persist under constant conditions, respond to timing cues, and display a temperature-resistant cycle length of around 24 hours^16^. These rhythms have been observed in every kingdom of life and across multiple scales of biological complexity, implying that they evolved to optimise survival on a planet that spins once daily^1,17^. Circadian ‘clocks’ thus broadly exemplify feedforward control, timing our physiology so that it crudely aligns with anticipated environmental change— these clocks predict time as much as they ‘keep’ it. Mechanistically, however, the oscillatory nature of molecular circadian phenomena is thought to be driven by delayed negative feedback^15^. It remains unclear as to whether these rhythms oscillate around an unachievable (or undesirable) homeostatic set point, whether circadian rhythmic amplitude represents the gain of an imperfect control system, or indeed whether this amplitude holds any biological significance at all. More important, perhaps, is that circadian clocks are imprecise but responsive; this adaptive ‘room for improvement’ means that actual external variation can fine-tune our internal clocks, keeping them in synch with the outside world^18^. Loss of circadian rhythm, or circadian rhythmic disruption, has been associated with negative health outcomes in both acute and chronic disease^9,19–23^. It is becoming increasingly clear that daily rhythms are important for brain health^2,9^ and, whilst causal links remain unproven, several neurodegenerative disorders feature disrupted daily rhythms years before clinical onset^23^. The demand for model systems in which human neural circadian phenomena can be measured and manipulated is therefore urgent and rising.

Our understanding of how circadian clocks operate extends from molecules to whole organisms, using an ever-expanding array of model systems that, with few exceptions, exploit the transcription-translation feedback loop (TTFL) put forward by Hall, Rosbash, and Young^15^. In this model, heterodimeric transcription factors (e.g. BMAL1 with NPAS2 or CLOCK) drive the expression of proteins (e.g. PER and CRY) that form cytoplasmic complexes; after sufficient build-up that takes several hours, these complexes traverse back into the nucleus to inhibit their own transcription by binding to their cognate transcription factors (**Fig.1a**). The cycle restarts once PER and CRY levels have fallen sufficiently to relieve this inhibition. Other TTFLs, along with several accessory loops, have since been described in a diverse range of organisms, suggesting that genetically distinct, yet functionally similar ‘cogs’ turn a ubiquitous circadian clock^24^. Whilst the absolute requirement of a TTFL for circadian rhythmicity remains disputed^25^, it has become the workhorse of chronobiologists seeking to observe circadian function in the lab. Typically, a bioluminescent or fluorescent reporter is tagged to a TTFL component so that circadian parameters (period length, phase, and amplitude) can be recorded under constant conditions in cells, tissue slices, or whole animals^26^. *In vivo* experiments have further targeted TTFL components in specific neural cell types, most notably in the suprachiasmatic nucleus (SCN) of the hypothalamus^27^. The rationale for these approaches is based on the putative hierarchical assembly of the circadian system in mammals. In this paradigm, the SCN ‘master pacemaker’ receives light signals via the retina and sets the pace of cellular circadian rhythms in the rest of the brain and body via neural, endocrine, thermal, and behavioural signals^28–29^.

**Fig. 1.**
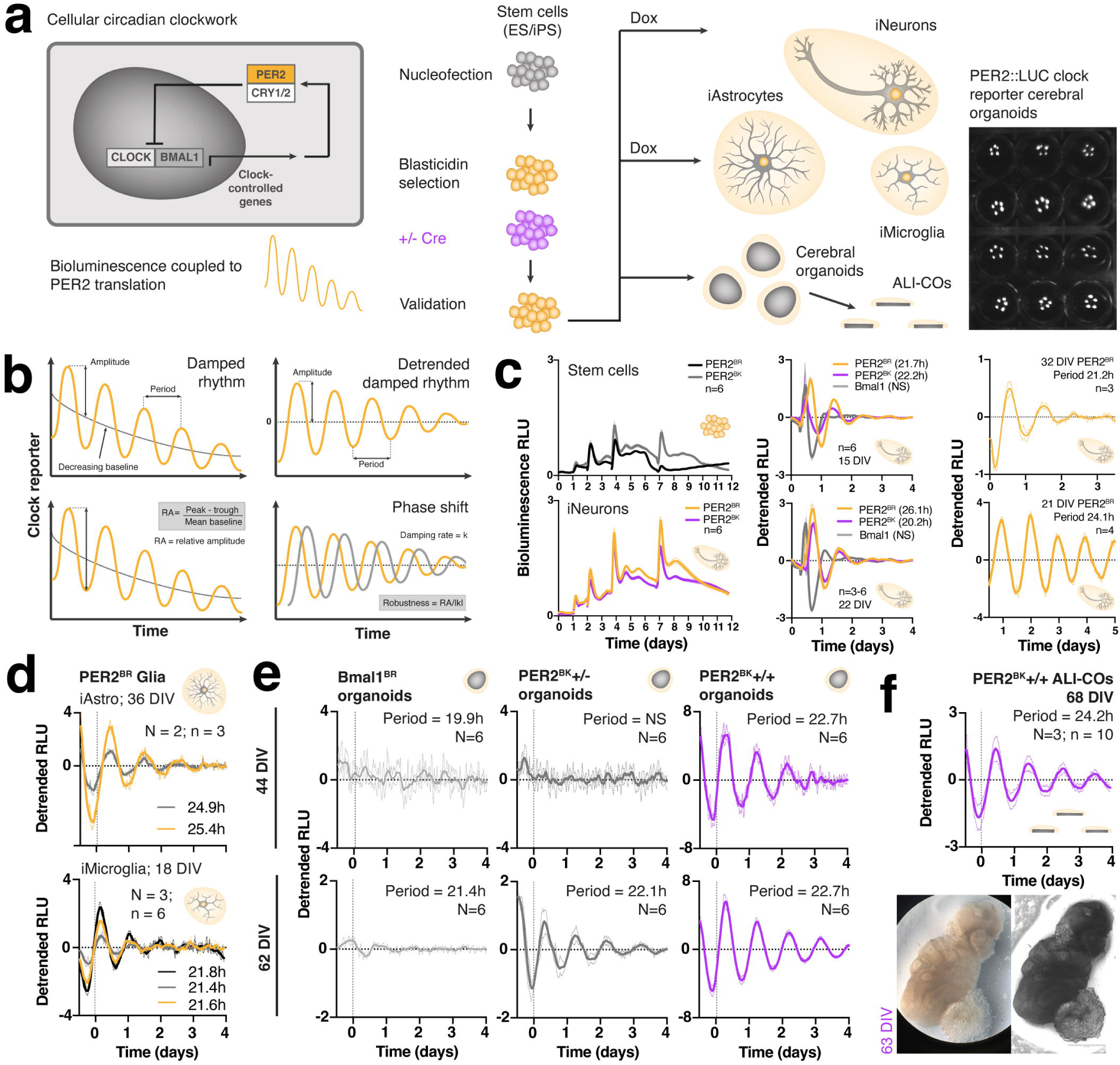
Circadian clock reporter human neural cultures. **(a)** Simplified depiction of molecular components of TTFL model in mammalian cells (left) exploited to create bioluminescent clock reporter human stem cells (middle). Clock reporter stem cells were differentiated into a range of neural types as well as organoids; far right image shows PER2+/+ organoids (N=60), with bioluminescence captured in ALLIGATOR (cultured at 5 organoids per well/condition; note spacing such that ROIs can be drawn around individual organoids for data extraction). **(b)** Schematic of extracted and derived parameters of a circadian oscillation. **(c)** Bioluminescent rhythms with metrics in PER2::LUC and *Bmal1*:Luc iNeuron platform (synchronized by media change). Left panel = changes in bioluminescence during differentiation comparing blasticidin-resistant (BR) and blasticidin-knockout (BK) biallelic knockin (PER2+/+) clones as stem cells versus iNeurons at 37°C. Broken curves reflect media changes on D1, D2, D4, D7. Note daily oscillations in iNeurons from D4 that rapidly dampen. Baseline stabilises from around D7, consistent with no further proliferation of cells. Apparent oscillations in stem cell cultures likely relate to spontaneous fibroblastic differentiation at point of overgrowth with rapid decrease in bioluminescence at D6 as many cells die off before D7 feeding. Data presented as mean of n=6 wells +/- 95% CI. Middle panel = PER2+/+ and *Bmal1* reporter iNeurons cultured to 15 or 22 DIV at 37°C before 30-40% minimal media change and acute shift to 32°C. Period lengths shown in brackets. Note rapid damping in all clones, lack of circadian rhythm in *Bmal1* iNeurons, and distinct phasing of *Bmal1* and PER2 initially. Top right = PER2^BR^+/+ iNeurons after 25% minimal medium change at constant 37°C, n=3 +/- SEM. Bottom right = PER2^BR^+/+ iNeurons cultured at 37°C then shifted to 32°C at zero days with 30% standard medium change; a further temperature drop to 30°C at 3.5 days (incubator malfunction) was associated with a reduction in relative amplitude. **(d)** Bioluminescent rhythms with metrics in PER2 iAstrocytes and iMicroglia (synchronized with glucocorticoid-containing media change and/or 100nM hydrocortisone). Note relative phase alignment of both culture types (6-8 h before the peak of PER2) in response to the same cue; time zero on the x-axis marks the start of the analysis window (24h after media change and into constant conditions). **(e)** Bioluminescent rhythms with metrics in cerebral organoids at D44 versus D62 with different reporters (all synchronized with 100nM hydrocortisone). From left to right: *Bmal1*, monoallelic (PER2^BK^+/-), biallelic (PER2^BK^+/+) organoids. Time zero is 24h after synchronization; note that all rhythms are set to same phase (GCT 0), consistent with molecular map of cellular circadian time developed in human and mouse fibroblast lines^37^. **(f)** ALI-COs from PER2^BK^+/+ organoids on inserts that underwent 100% medium change followed by re-synchronization with 100nM hydrocortisone 11h later. Time zero is 24h after glucocorticoid was added. Note perfect phase alignment with PER2+/+ organoids in **(e)**. Brightfield and phase-contrast images of 63 DIV ALI-CO sliced from PER2^BK^+/+ organoid shown in lower panel.

In rodents, the hypothalamic circuitry that coordinates cycles of rest and activity with those of day and night is well characterized^27^. Far fewer studies have explored circadian phenomena in other parts of the brain^30–33^, and fewer still have addressed them in naturally diurnal species. This has proven to be technically challenging, leading some to propose that neural circadian rhythms are generally weak or absent beyond core, diencephalic structures^30^. Such conclusions have often relied on the use of a transcriptional clock reporter, and it has since been shown that transcriptional and translational circadian rhythms may be poorly correlated^34^. Notwithstanding that rhythms in a clock reporter may not reflect direct observation of the clock itself, and that interpretation of these rhythms must consider the kinetics of the readout^35^, many useful insights can be gleaned from TTFL-based bioluminescent platforms. For example, the effects of temperature on circadian rhythms *in vitro* are markedly different in mouse and human cells, whilst other key timing cues set them to the same circadian phase^36–37^. This is paramount given the distinct temporal niches of these species, their differing thermoregulatory profiles, and that nocturnal rodents are the mainstay of preclinical studies, often with poor translation to humans^38–39^.

*Bona fide* rhythms that meet established circadian criteria have not yet been demonstrated in the human brain or any human neural platform. To address these gaps, and to overcome inherent limitations of overexpression systems, our primary goal was to tag endogenous human PER2 with firefly luciferase^36^, generating a translational reporter of circadian clock function (PER2::LUC) through CRISPR/Cas9 gene editing of human pluripotent stem cells (**Fig.1a and Extended Data Fig.1a, Supplementary Fig.1a**). In parallel, we took the more conventional approach of generating transcriptional clock reporter stem cells (*Bmal1*:Luc) via lentiviral transduction^40^. Exploiting stem cell lines that already carry the potential for doxycycline-induced directed differentiation by cell reprogramming via opti-ox™ technology^4–5^, we were able to establish bioluminescent clock reporter clones that could be differentiated into one of three key brain cell types or, in the absence of doxycycline, developed into cerebral organoids (**Fig.1a**)^7^. Using orthogonal approaches including bioluminescence, mass spectrometry, and electrophysiology, we show that human brain cells and tissues exhibit true circadian oscillations. In contrast to *ex vivo* platforms, our work establishes that circadian rhythms can develop in brain cells devoid of any historical influence from a functioning hypothalamus, or light-receptive circuitry. This fundamentally challenges the developmental origin of circadian timekeeping in mammals^41–42^, and provides resources with which to explore human neural circadian rhythms under physiologically-relevant conditions, as well as in different contexts relevant to brain health.

### Circadian rhythms in human brain cells and organoids

To generate clock reporter human stem cells efficiently, we exploited antibiotic resistance to select for edited or transduced clones. Blasticidin was chosen since the opti-ox™ technology renders stem cells resistant to both puromycin and neomycin, and blasticidin effected rapid cell death of small colonies. A floxed resistance cassette was included in our editing strategy (**Extended Data Fig.1a**); this was subsequently removed by a cell-permeant cre-recombinase to test for any effect of blasticidin resistance on circadian parameters (**Fig.1b**). Bioluminescence was recorded under constant conditions in an ALLIGATOR system (Cairn Research)^43^ (**Fig.1a, Extended Data Fig.1b**). No significant differences were observed in the circadian bioluminescence of blasticidin-resistant (BR) versus isogenic blasticidin-knockout (BK) clones, either before or after differentiation (**Fig.1c-f**). By editing different opti-ox™ stem cell lines we generated highly enriched bioluminescent clock reporter cultures of glutamatergic excitatory cortical neurons (iNeurons^4^, **Supplementary Video 1**), astrocytes (iAstrocytes^5^, **Extended Data Fig.2, Supplementary Fig.1b**), and microglia (iMicroglia, using the <u>ioMicroglia™</u> model from bit.bio Ltd). Excluding doxycycline from our stem cell cultures enabled generation of isogenic bioluminescent cerebral organoids^7^ (**Supplementary Fig.1c, Supplementary Video 2**), from which air-liquid interface cerebral organoids (ALI-COs^8^, **Fig.1f**) were sliced. For comparison, we transduced opti-ox™ iNeuron stem cells and previously characterized astrocyte neural progenitors^6^ with a *Bmal1*:Luc transcriptional reporter^40^. Under constant conditions, neural rhythms in *Bmal1*:Luc were generally poor, rapidly damping, or absent (**Fig.1c and e, Extended Data Fig.1c, f-g**). By contrast, PER2::LUC rhythms were either adequate (monoallelic knockin, PER2+/-) or excellent (biallelic knockin, PER2+/+ **Fig.1c-f, Extended Data Fig.1c-h**), enabling reliable extraction of circadian parameters. Notably, PER2::LUC rhythms became more robust at later stages of differentiation, with period lengths between 21 and 27 hours, depending on culture type and age (**Fig.1c-f, Extended Data Fig.1c-h**). iNeuron rhythms were less reproducible than other platforms, glial rhythms were phase-advanced by a few hours, and PER2::LUC rhythms were most persistent in organoids (**Fig.1c-f, Extended Data Fig.1c-h**). The peak of reported PER2 translation was almost antiphasic with that of *Bmal1* transcription (where this could be discerned) and bioluminescence rhythms were readily observed after synchronizing with conventional timing cues. Glucocorticoid synchronization produced notably high-amplitude rhythms, setting circadian phase in organoids and ALI-COs to our previously defined reference of glucocorticoid time zero (GCT 0; approximately 10h before the peak of PER2::LUC and 19h prior to the peak of *Bmal1*:Luc)^37^. These data demonstrate that, similar to other mammalian cultures, human neural platforms display daily rhythms in bioluminescence, coupled to the transcription and translation of key TTFL components.

### Human neural clocks are sensitive and resistant to temperature

To test for temperature compensation of period length (**Fig.2a**), human neural clock reporter cultures were incubated at different constant temperatures during bioluminescence recording. This included 37°C and 40°C (spanning the range of normal human brain temperatures^9^) as well as 32°C–reflecting the core body temperature approximated during moderate therapeutic hypothermia in critical care^44^. Cultures readily tolerated 32-40°C (**Supplementary Fig.2a**) and period length was stable across the physiological range (**Fig.2a, Extended Data Fig.3a**). There was a trend for period lengthening in organoids as temperature increased, and period shortening as organoids aged, or if they were synchronized with stronger timing cues (feeding versus temperature cycling) (**Fig.2a**). An array of acute temperature upshifts and downshifts were applied to demonstrate temperature-driven circadian phase resetting (**Fig.2b**). Across different neural culture types, temperature increases generally set rhythms to opposing phases relative to temperature decreases^36–37^. Astrocyte rhythms were consistently phase-advanced by around 4 hours relative to other culture types, and the presence of non-bioluminescent iAstrocytes modified the sensitivity of PER2::LUC to temperature cycles in co-cultured clock reporter iNeurons (**Fig.2b, Extended Data Fig.3b-c**). The advanced timing of glial rhythms was reproducibly observed in progenitor-derived human astrocytes^6^, even though antiphasic temperature cycles predictably entrained these monocultures to opposing phases (**Extended Data Fig.3b-e**). Organoid rhythms were more robust at physiological brain temperatures, below which *Bmal1*:LUC (but not PER2::LUC) rhythms were notably absent (**Extended Data Fig.3d, Supplementary Fig.2b**). To confirm whether physiological human brain temperature cycles^9^ can reset the phase of neural circadian rhythms, we performed a temperature re-entrainment experiment with organoids (**Fig.2c**). Applying a conservative 39/37°C cycle over 4 days was sufficient to re-align desynchronized organoid rhythms into the same circadian phase (**Fig.2d-e; Extended Data Fig.3f-g**). A similar result was achieved with iAstrocytes and ALI-COs, noting again that iAstrocyte rhythms were phase-advanced by 4-6 hours compared to organoid and ALI-CO rhythms (**Fig.2f**). Thus, human neural bioluminescence rhythms meet hallmark temperature criteria for circadian rhythmicity: phase resetting to temperature shift, and temperature compensation of period length.

**Fig. 2.**
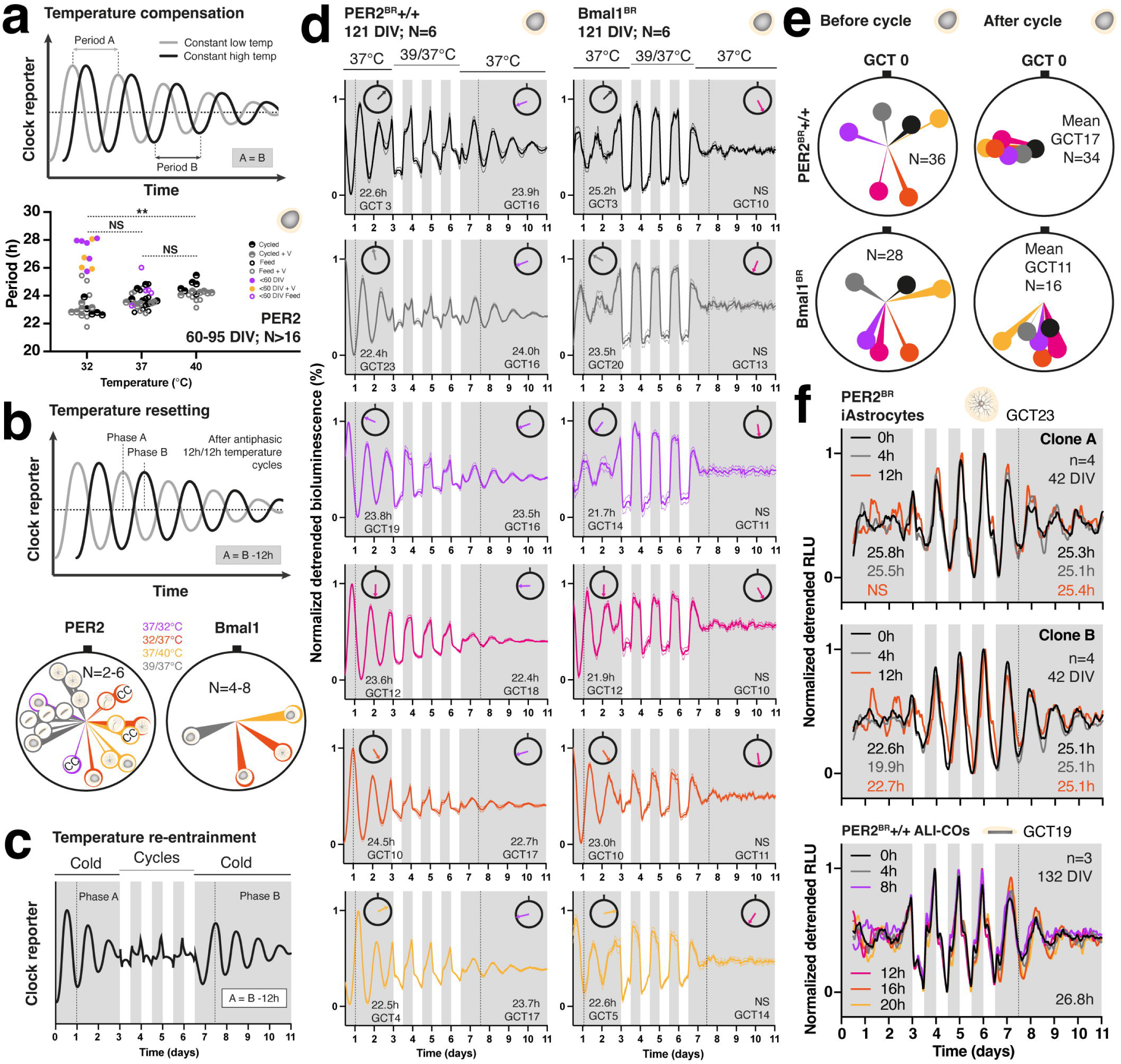
Temperature compensated neural circadian clocks entrain to human brain temperature cycles. **(a)** Schematic outlining temperature compensation of circadian period length (top). Graph (bottom) shows confirmatory data for PER2+/+ organoids under constant conditions at different temperatures, having previously been temperature cycled or not, fed only, or fed with vehicle, and at different ages in culture. Note lack of difference in period length across the physiological range of human brain temperatures (37-40°C)^9^, with variable period length at sub-physiological temperature of 32°C that is significantly shorter than at 40°C in older organoids (statistical test applied for organoids 60+ DIV only as 40°C data not available for younger organoids; **p = 0.002; Kruskal-Wallis with Dunn’s correction for multiple comparisons). **(b)** Schematic demonstrating temperature resetting. Circular graphs in bottom panel show how reporter peaks are set by different temperature shifts (colour-coded) in various types of human neural culture, depicted by same cartoons used in Fig.1. CC=iNeuron/iAstrocyte co-culture in which bioluminescent iNeuron rhythms were recorded when co-cultured with non-bioluminescent iAstrocytes. Each large circle represents 24 circadian hours; note that PER2 peak ends up on roughly opposite sides of the circle when shifted up in temperature relative to down, and that the peak of *Bmal1* occurs at a very different time relative to PER2 as expected. Width of each wedge corresponds to circular SD. **(c)** Paradigm of temperature re-entrainment experiment in organoids, predicting that circadian rhythms in different phases to each other can be shifted to the same phase using a simulated brain temperature cycle. **(d)** Confirmatory bioluminescence data for **(c)**; 6 batches of N=6 organoids were synchronized by full media change with additional 100nM hydrocortisone at 4-hourly intervals to misalign them with respect to external time ahead of 12h/12h temperature cycling (39/37°C). Note that 4 cycles is sufficient to set all rhythms to the same final phase. Note also that *Bmal1* bioluminescence responds well initially to the glucocorticoid stimulus but *Bmal1* rhythms after the temperature cycle are poorly discernible and grouped data fail to reach significance for a cosinor fit; interpretation of the experiment is best made with PER2. Mini clocks highlight the phase of the rhythm (in circadian hours, thus normalizing for variable period lengths) before and after the temperature cycle, to the nearest circadian hour, relative to GCT 0 (black notch). **(e)** Circular quantification of **(d)** highlighting re-synchronization of organoids after the temperature cycle. Width of each wedge corresponds to circular SD. **(f)** Parallel temperature re-entrainment data collected from iAstrocytes and ALI-COs; note that ALI-CO rhythms end up in roughly the same phase as organoid rhythms after the temperature cycle, whilst iAstrocyte rhythms are phase-advanced (compare dotted vertical lines and GCT values). Two independent clones of iAstrocytes (A and B) were tested across 3 initial phases; one ALI-CO clone was tested across 6 initial phases. In **(d)** and **(f)**, period lengths are displayed in hours where the cosinor fit was significantly preferred over a straight line for grouped data (otherwise NS) and mean GCT values of the resultant phase are given to the nearest circadian hour, whilst the x-axis displays external time. Vertical dotted lines signify phase of rhythm 24h after the start of constant conditions. For Bmal1 organoids, the quantification of phase (and thus GCT) included only individual organoids whose bioluminescence was significantly rhythmic. Colour coding corresponds to different phasing before the temperature cycle. Normalization of detrended data refers to normalization for amplitude during each stage of the experiment (before, during, and after temperature cycling).

### Development of organoid clock responses to glucocorticoid

In the adult brain, glucocorticoid receptors (GRs) are widely expressed but have a 10-fold lower affinity for glucocorticoid relative to mineralocorticoid receptors (MRs), whose expression is greater in certain regions like the hippocampus^45^. Whilst glucocorticoids bind to both receptor types, brain sensing of daily rhythms in cortisol is thought to be mediated via GRs, since MRs are fully occupied at trough concentrations of cortisol, whilst GRs only become activated at peak levels^46^. During neurodevelopment, however, there is a transition in the functional activity of GRs and MRs in the brain such that GR affinity for glucocorticoid is high initially, but reduces as MR expression increases^47^. We used our recently-established molecular timetable of cellular circadian phase^37^ alongside GR- and MR-selective antagonists to explore glucocorticoid-induced phase resetting of organoid rhythms at various stages of differentiation^48^. In mouse and human fibroblasts, and in the absence of other timing cues, glucocorticoid reliably sets circadian phase to GCT 0. Growth factors such as insulin set rhythms to a later phase, consistent with the temporal order of circadian physiology (**Fig.3a**)^37^. Cell culture supplements that contain both glucocorticoid and insulin (e.g. B-27), set cells to around GCT 2, reflecting the resultant phase of conflicting cues, but also the greater strength of glucocorticoid relative to insulin^37^. Since organoid culture media contains B-27 and extra insulin, we reasoned that adding more glucocorticoid with media replacement would set rhythms to around GCT 0-2 and improve rhythmic amplitude via enhanced synchronicity. We further hypothesised that the greatest phase shift would be achieved if glucocorticoid was applied at around GCT 12 (the most ‘incorrect’ biological time), but that this response may (a) take time to develop in culture and (b) depend on glucocorticoid type^49^. Accordingly, adding hydrocortisone at GCT 12 reduced the variability in period length of individual organoids, and increased the relative amplitude of rhythms beyond that achieved by ageing in culture alone (**Fig.3b, Extended Data Fig.4**). Robustness was also enhanced in an age-dependent manner, but more so in the presence of added glucocorticoid. Phase-shift in response to glucocorticoid was only notable from D38, being greatest with hydrocortisone, a synthetic mimic of the principal endogenous human glucocorticoid (cortisol) (**Fig.3c, Extended Data Fig.4**). At this stage, a smaller phase shift was noted with the highly potent, GR-specific synthetic glucocorticoid dexamethasone, but corticosterone was entirely ineffective. By D44, a full phase resetting was achieved with all 3 glucocorticoids. The response was somewhat variable up to D71 but remained optimal using hydrocortisone at this later stage. Of note, the MR-selective antagonist spironolactone (S) showed an age-dependent partial blocking effect on hydrocortisone resetting (**Fig.3c, Extended Data Fig.4**). By contrast, the GR-selective antagonist, exicorilant (X, CORT125281, Corcept Therapeutics)^48^ effectively blocked resetting by all 3 glucocorticoids at most time points, but was specifically less effective at abrogating hydrocortisone resetting after organoids had been cultured at 32°C. Importantly, S reduced the relative amplitude of rhythms at all stages, preventing the enhancement achieved by hydrocortisone except at the latest time point (D71), whilst X had the opposite effect (**Fig.3c, Extended Data Fig.4**). Moreover, the damping effect of S at D71 was phase-dependent and, at this latest time point, the impact of glucocorticoid on both phase shift and amplitude was greatest for hydrocortisone > dexamethasone > corticosterone. In summary, organoid circadian phase resetting by glucocorticoid is rapid (<30min) and best achieved with hydrocortisone; this resetting is robust by D44, with amplitude enhancement taking longer to develop. Whilst GRs likely play the dominant role in neural phase-resetting, our data suggest that MRs contribute to this and appear to be the primary mediators of elevated amplitude in response to hydrocortisone. Our results recapitulate the relative affinities of the MR and GR for glucocorticoid, and developmental transitions in the activity of MRs versus GRs in the brain^11–14,45–47^.

**Fig. 3.**
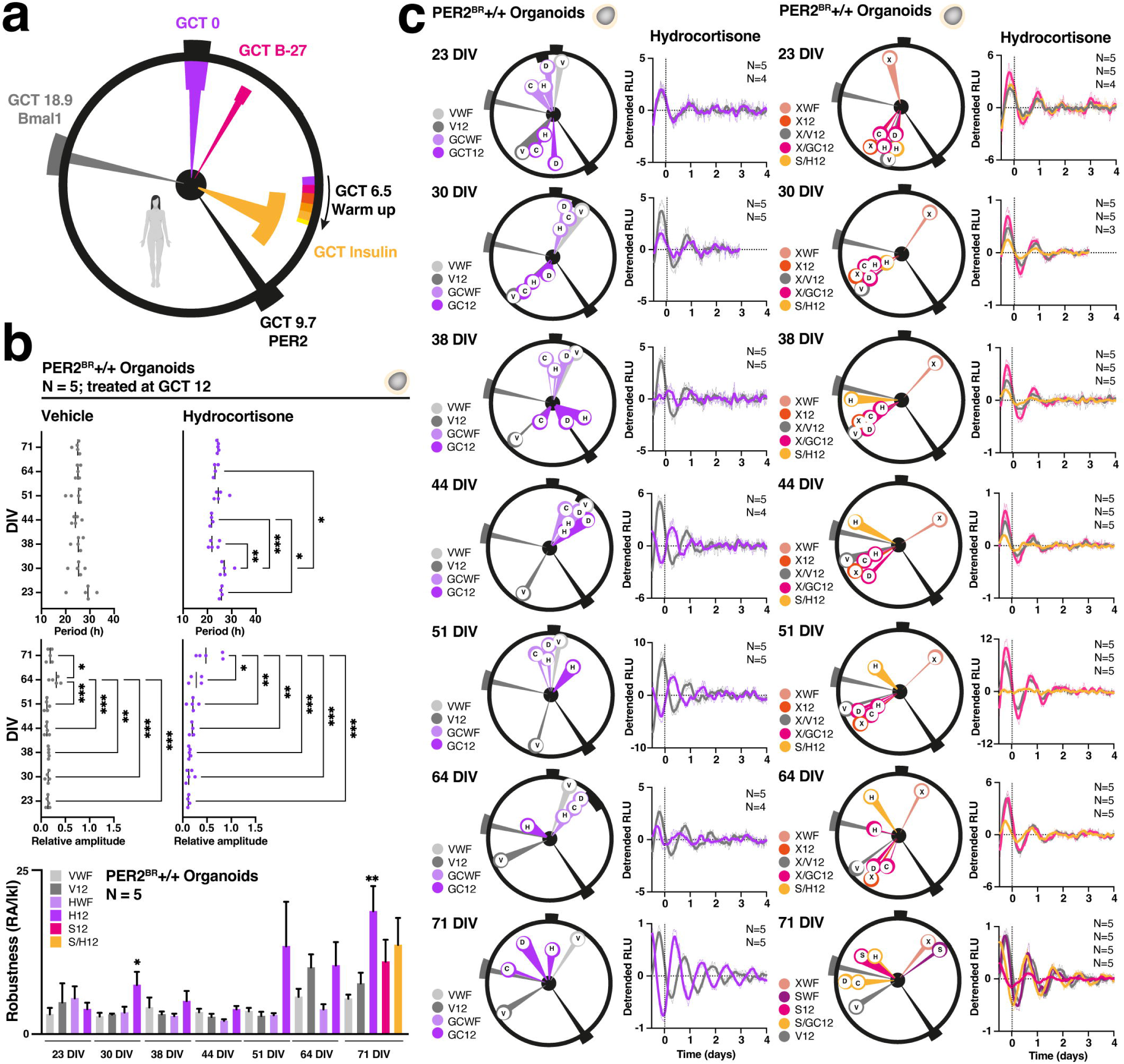
Organoid clock responses to glucocorticoid reflect neurodevelopmental transitions. **(a)** Key timing cues and the circadian phases they set cells to, based on a molecular map developed in 15 independent human and mouse fibroblast lines^37^; the large circle presents 24 circadian hours. Specific phases are depicted relative to GCT 0, defined as the phase cells are set to by glucocorticoid. The peaks of the PER2 translational reporter (black) and *Bmal1* transcriptional reporter (grey) are shown at their respective GCTs. GCT of insulin (orange) and B-27 (magenta) are shown since these are included in the standard culture media for organoids which was used for all feeds prior to experimental runs. Note that the phase setting effected by B-27 lies between GCT 0 and GCT insulin but nearer to GCT 0, since B-27 contains both corticosterone and insulin, and glucocorticoid is the stronger timing cue. The phase set by a temperature increase is relevant to human cell lines only^36–37^ and is consistent with data shown in Fig.2. **(b)** Period, relative amplitude, and robustness data from bioluminescence rhythms recorded in PER2^BR^+/+ organoids at various ages in culture from 23 to 71 DIV. Note progressive increase in relative amplitude and robustness over time, with a period length that stabilises and becomes more reproducible from around D38. Relative amplitude and robustness are increased further with the addition of hydrocortisone given at GCT 12, most apparent at D71. GCT 12 was timed to effect the greatest phase-shift based on (a), assuming that 100% media change would set rhythms to around GCT 2 (GCT B-27). Therefore, glucocorticoid (or DMSO vehicle) was applied 10h after media change. The robustness graph also features parallel effects of applying hydrocortisone with media replacement (WF) versus GCT 12 as well as the effect of 10uM spironolactone (S). For double-treated organoids, S was applied at GCT 11.5 (30min prior to hydrocortisone at GCT 12). One-way ANOVA with Tukey’s correction for multiple comparisons, *p<0.05, **p<0.01, ***p<0.001 are displayed in separate graphs. Mixed effects analyses with correction for multiple comparisons highlighted significantly shorter period length for hydrocortisone relative to vehicle at D44 (p=0.02) and D64 (p=0.01), and a shorter period length for hydrocortisone at D44 versus D71 (p=0.03). Relative amplitude was greater for hydrocortisone compared to vehicle at D44 and D71 (both p=0.03). **(c)** Circular quantification of phase responses to glucocorticoid and steroid antagonists applied with media change (WF) or at predicted GCT 12. Key synthetic and natural glucocorticoids were tested in parallel. V (DMSO vehicle at final concentration of 0.1%), D (100nM dexamethasone), C (100nM corticosterone), H (100nM hydrocortisone). Inhibitors included a selective glucocorticoid receptor (GR) antagonist exicorilant (X, CORT125281, Corcept Therapeutics, Menlo Park, CA, USA)^48^ at 10uM or spironolactone (S) at 10uM, applied WF, at GCT 12, or GCT 11.5. Each wedge represents the resultant circadian phase relative to GCT 0; width of each wedge refers to circular SD and predicted peaks of PER2 and *Bmal1* reporters are shown. Black tags internal to the large circle reflect ‘untreated’ result i.e. 100% media change alone– note that by D44 this achieves the predicted phase of GCT 2 (B-27). Left panel: note that glucocorticoid resetting commences by D38 (H>D>C), with resetting to GCT 2 achieved for H and D by D44. Only data for H were available at 51 DIV (resets to GCT 2) and at 64 DIV (resets to GCT 20) which was recorded after one week at 32°C (56-63 DIV) with a shift back to 37°C at the point of media change; this affected the resetting capacity of H. At 71 DIV, H achieves full resetting to GCT 0 and over-rides any effect of other components in the media. Representative bioluminescence traces for each developmental stage are shown for hydrocortisone given at GCT 12. Right panel shows effects of steroid antagonists which initially have little to no impact on rhythmic timing. From D38, X completely, and S partially, abrogates the resetting effect of H. Representative bioluminescence traces of key results are shown, with x-axis plotting in standard time, and circular plots in circadian time (normalized for period length). Vertical dotted lines at 0 days marks the start of the analysis window (24h after start of constant conditions). The absence of data for a cue at any given time point reflects loss of that well of organoids due to fungal contamination in the ALLIGATOR system.

### Multiomics of circadian organoid clocks

To fully exclude the possibility of a bioluminescent reporter phenomenon, we sought to extract a more comprehensive analysis of circadian physiology in cerebral organoids. Taking a multiplexed, multiomics approach, we designed a time course to collect several readouts in parallel from D62 organoids over 2 days (**Fig.4a**). This culture age was chosen based on our earlier data (**Fig.3**), ensuring robust synchronization of organoids with hydrocortisone at experiment start. The time course included two independent batches of PER2^BK^+/+ organoids maintained at constant 38.5°C (mean global brain temperature in healthy adults)^9^. Bioluminescence confirmed that organoid PER2::LUC rhythms were set to GCT 0 at experiment start (**Fig.4b, Extended Data Fig.5a**), and temperature-controlled sampling commenced 24h later. Protein samples (N=3 organoids per sample) were collected every 3 hours enabling 18-plex TMT mass spectrometry with a temporal resolution of 8 circadian phases. We detected 9008 proteins, with 7187 (80%) detected at all time points in both batches, and 50,532 phosphopeptides with phosphosite localization probability >0.75 (mean 0.97^50^), mapping to 5051 unique proteins (93% of which were captured in the total proteome). 9431 (19%) of the phosphopeptides were detected in all samples and 8245 (87%) of these derived from proteins that matched Uniprot IDs in the ‘all samples’ proteome (**Fig.4c**). These filtered phosphopeptides represented 7305 unique phosphosites mapping to 2640 proteins; we noted similar proportions of phosphorylated amino acids as described before^50^, varying slightly by multiplicity (**Supplementary Fig.4a-c**). Only proteins and phosphopeptides that were detected in all samples (at all time points) were included in circadian rhythmicity analysis by RAIN^51–52^. Since 912 (35%) of the 2640 phosphoproteins were represented in the RAIN-defined rhythmic proteome, phosphopeptide abundance was normalized to respective protein abundance prior to RAIN analysis^52^. On this basis, 33% of the proteome and 27% of the phosphoproteome was rhythmic, and rhythmic proteins and phosphopeptides had a significantly greater abundance (**Fig.4d, Extended Data Fig.5b**). The rhythmic proteome was notably enriched for ribosomal RNA processing, protein targeting to the membrane and endoplasmic reticulum, and synaptic processes (**Fig.4e, Extended Data Fig.5c-d**). Accordingly, rhythmic molecular functions included RNA binding and structural constituents of the ribosome, whilst rhythmic cellular components were enriched for ribosomal subunits, the synapse, and axonal cytoskeleton. The rhythmic phosphoproteome was primarily enriched for targets involved in neurogenesis and actin dynamics (**Extended Data Fig.5c-d**)^29^. Next, we clustered the rhythmic hits by phase of peak abundance to generate a ‘circadian organoid multiomics clock’ (**Fig.4f**). This reassuringly highlighted enrichment for the cortisol response at GCT 0, RNA splicing and DNA repair at GCT 3, protein turnover at GCT 6, microtubule regulation at GCT 12 and synaptic signalling at GCT 12-15. Most rhythmic phosphopeptides peaked in the early part of the circadian cycle, whilst rhythmic proteins were more widely distributed, but GCT 21 was distinctly barren for both. We noted that the several epitopes of disease-relevant microtubule-associated protein tau (MAPT) that were rhythmically phosphorylated peaked in their phosphorylation at the opposite phase to MAPT abundance, with biomarker NF-L (NEFL) peaking 3 hours later (**Extended Data Fig.5d**). Finally, we noted that the relative amplitude of rhythmic proteins and phosphopeptides varied by phase of peak abundance (**Fig.4g**); moreover, relative amplitude was greater at the post-translational level in most phases of the circadian cycle^50,52–53^, with the exception of GCT15-18. Thus, we present circadian mass spectrometry datasets from human cerebral organoids, identifying rhythmic features of the neural cellular machinery that very likely impact on diurnal brain function.

**Fig. 4.**
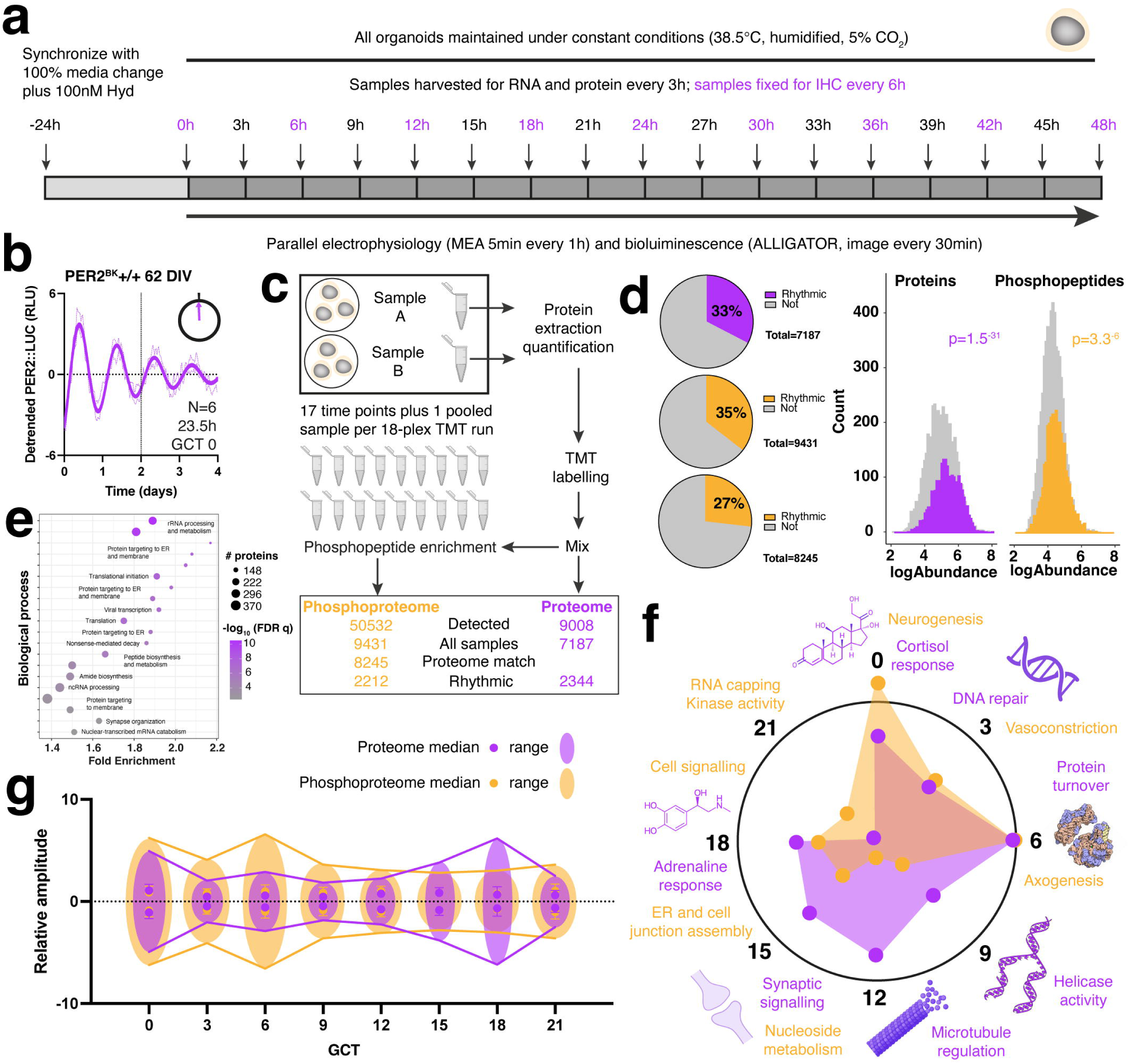
Building a circadian multiomics clock in human cerebral organoids. **(a)** Schematic of multiomics paradigm. D62 PER2^BK^+/+ organoids were synchronized with 100% media change plus 100nM hydrocortisone and maintained under constant conditions. N=6 organoids were immediately transferred to an ALLIGATOR for bioluminescence recording, and another N=6 underwent MEA recording on the Axion Maestro Pro. Regardless of incubation device, all organoids were maintained at 38.5°C throughout the time course (or until harvested). N=3 organoids were fixed every 6h for IHC. N=6 organoids were harvested very 3h for multiple RNA readouts, and N=6 organoids were harvested every 3h for protein readouts (2 independent batches of 3 organoids were collated into 2 samples (A and B) per time point). **(b)** Bioluminescence quantification showing nonlinear fit of grouped data +/- SEM of unsmoothed detrended raw data. Period length is shown. Time zero on the x-axis refers to 24h after synchronization (when sample harvesting commenced) and is phased at GCT 0 (∼10h before the peak of PER2 translation). Vertical dotted line denotes end of sample harvesting window. **(c)** Schematic and flow chart of protein sample analysis by mass spectrometry; there were 17 time points for each TMT run (A and B); the 18^th^ sample for both runs was an identical composite of equal amounts of all 34 samples, used as an internal reference. **(d)** Pie charts showing proportions of rhythmic proteome and phosphoproteome together with frequency histograms highlighting greater abundance of rhythmic (coloured) versus non-rhythmic (greyscale) proteins and phosphopeptides. Middle pie chart shows proportion of phosphoproteome that was rhythmic before phosphopeptides were normalized to protein abundance. P-values refer to t tests comparing abundance of rhythmic versus non-rhythmic proteins and phosphopeptides, performed using log- transformed data. **(e)** Bubble enrichment plot for rhythmic proteome showing top 20 biological processes according to GO analysis fold-enrichment and FDR q-value. Four rhythmic proteins (LANCL1, STMN2, ALDOC, and NRXN3) were recently identified as correlates of human brain ageing^96^ and another (PCLO) in neuropsychiatric disorders^97^. **(f)** Circadian organoid multiomics clock comprising 8-point radar chart for phase-clustered rhythmic proteome (purple) and phosphoproteome (orange). Each radial point plots the log2 proportion of rhythmic hits having their acrophase of abundance at each circadian phase. Terms surrounding the clock highlight the most prominent biological processes according to GO analysis; note temporal consolidation. **(g)** Forth Bridge (cantilever) plot showing relative amplitude of rhythmic hits by phase, with protein and phosphopeptide data overlaid. Median +/- SD and range are duplicated above and below y=0 to avoid inference of directionality, which is irrelevant in this context. Relative amplitude differed significantly between proteins and phosphopeptides (p=0.012, Kruskal-Wallis test with Dunn’s correction for multiple comparisons). Schematics created with BioRender.com, DNA replication schematic modified from ‘DNA replication split.svg’ by Madeline Price Ball under CC BY-SA 3.0 license, ribosomal subunits generated by David S Goodsell and downloaded from http://doi.org/10.2210/rcsb_pdb/mom_2000_10.

### Human neural activity around the clock

Our mass spectrometry data highlighted circadian rhythmicity in several biological processes and cellular components related to functional neural activity. To confirm this, we exploited multielectrode array (MEA) platforms to measure extracellular field potentials in 2D and 3D neural cultures around the clock. For monolayers and organoids we used the Maestro Pro MEA system (Axion Biosystems) (**Fig.5a**), enabling temperature-programmable, longitudinal data collection over multiple days with the same constant conditions we had employed for other readouts. At D29, neuronal spikes and bursts were readily detectable in iNeurons, but at a higher frequency and with longer burst duration in the presence of iAstrocytes (**Fig.5b, Extended Data Fig.6**). Network bursts were slightly more frequent in early iNeuron monocultures but substantially more frequent in co-cultures at D74 (**Supplementary Fig.5**); mono- and co-cultures responded predictably to established excitatory and inhibitory neurotransmitters and ion channel modulators^54^. Importantly, the spiking and bursting of both culture types varied in a circadian fashion (**Fig.5c, Extended Data Fig.6a,c**), but bursting and network bursting rhythms displayed a higher relative amplitude in the presence of iAstrocytes, in agreement with the range of data across the entire time course (**Fig.5b**). Phase shifts in these rhythms were observed in the presence of hydrocortisone which had variable, age-dependent modulatory effects on electrophysiological parameters in iNeuron and co-cultures. Overall, we observed a general suppression of electrical activity alongside an increase in the synchronous firing of entire networks by hydrocortisone, particularly in the presence of iAstrocytes (**Fig.5b-c, Extended Data Fig.6a-c**). Since rhythmic functional activity was best appreciated with added hydrocortisone, this was included when recording from 3D cultures; we noted circadian variation in spikes and bursts in whole organoids using the Axion device (**Fig.5d, Extended Data Fig.6d**), and in ALI-COs, using a bespoke engineered device for tissue slices (**Fig.5e, Extended Data Fig.6e**). In summary, these data confirm that human neural cultures display circadian rhythms in functional activity that are modulated by astrocytes and glucocorticoid.

**Fig. 5.**
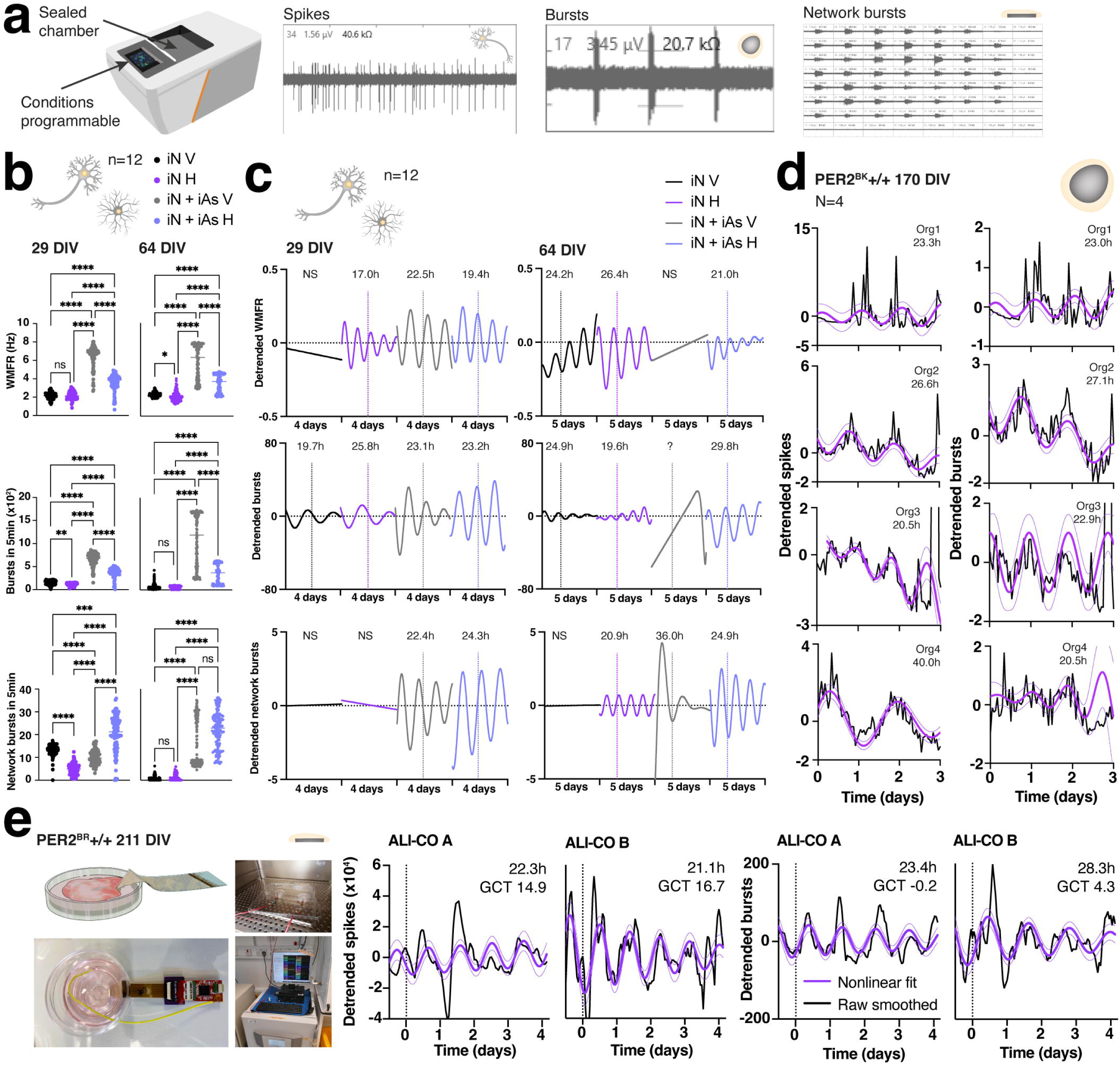
Circadian rhythms in human neural functional activity. **(a)** Maestro Pro MEA system (Axion BioSystems) device used for high-throughput MEA recordings with example images showing spikes (field potentials from an iNeuron monoculture), bursts of spikes in close succession (from whole organoids), and a network burst reflecting synchronous firing across the culture (from an ALI-CO slice). Maestro Pro MEA system image used with permission from Axion BioSystems Inc. **(b)** Summary MEA data from iNeuron monocultures and iNeuron-iAstrocyte co-cultures in a 48-well plate (n=12 wells per condition). Recordings of the same plate were made for 4 days from D29 and 5 days from D64, sampling 5 minutes’ worth of data every hour under constant conditions (humidified 5% CO_2_, 37°C). The weighted mean firing rate (WMFR), numbers of bursts, and number of network bursts recorded across the entire time course are shown. Treatments included with media change at start of recording were 0.1% DMSO (vehicle = V) and 100nM hydrocortisone (H). Horizontal bars represent median; all data points are shown. Asterisks report results of Kruskal-Wallis test with Dunn’s correction for multiple comparisons (*p<0.05, **p<0.01, ***p<0.001, **** p< 0.0001, ns = not significant). **(c)** Same data from **(b)** are presented as nonlinear fits of the time courses, staggered on the x-axis by condition for visualization. See Extended Data Fig.6 for raw data. Period lengths are displayed if the cosinor fit was preferred over a straight line; the analysis window commenced 1 day after the start of recording (true constant conditions) even though the displayed fit has been extended across the entire recording period. Vertical dotted lines at 2 days after the start of recording enable rough comparison of phases between conditions; the x-axis denotes standard rather than circadian time (not normalized for variations in period length). Note that circadian rhythms in network bursts of co-cultures are approximately antiphasic in V versus H conditions at D29, and a similar result is observed for WMFR of iNeuron monocultures at D64. **(d)** Circadian rhythms in spikes and bursts of N=4 whole organoids cultured at 37°C. Detrended (unsmoothed) data, nonlinear fits, and period lengths are shown for each organoid; note that best fit for Org4 suggests a doubling of period length for spikes relative to bursts although <24h variation can be appreciated in the underlying detrended data. **(e)** Circadian rhythmic data collected from two individual ALI-COs (A and B) using a bespoke engineered MEA device that enables long-term recordings from tissue slices in a standard incubator (humidified, 5% CO_2_, 37°C). A schematic and photos of the device and setup are shown, with one ALI-CO per device, and two ALI-COs recorded in parallel. Just prior to recording start, both ALI-COs underwent 100% media change with 100nM hydrocortisone added. Detrended spikes (in either direction) and bursts (positive direction) are presented for each ALI-CO (black traces) with nonlinear fit overlaid (purple). Vertical dotted lines at time zero represent the start of true constant conditions (24h after start of recording) from which point the cosinor analysis window commenced. Period lengths and phases (in GCT) are displayed on each graph; note that phases are similar between ALI-COs for a given readout, but differ between readouts. Nonlinear fits represent best approximation of underlying rhythm based on cosinor analysis (as is standard in the field); note that the true rhythms are non-sinusoidal and asymmetric as noted in mass spectrometry data (**Fig.4**).

### Circadian disruption in human brain clocks

Modern lifestyles and working schedules challenge our circadian system by repeatedly misaligning our internal clocks with external time^2^. Such circadian disruption has been cited as a major risk factor for chronic brain disorders from psychiatric illness to neurodegeneration and has been reviewed extensively elsewhere^55^. To demonstrate the disease-modelling value of our bioluminescent organoid platform, we subjected D68 PER2^BK^+/+ organoids to a ‘shift work’ paradigm (**Fig.6a**), exploiting that fact that, *in vivo*, tissue cortisol levels peak at around the time of waking^56^, preparing our physiology for the active phase. Batches of organoids were synchronized with 100% media change then, starting from 24h later, 100nM hydrocortisone was added to batches at 4-hourly intervals, constituting 6 phases over 24 hours. Bioluminescence was then recorded under constant conditions at 37°C. Since the glucocorticoid cue was being applied on a background of several other conflicting cues in the culture media, the extent of resetting was phase-dependent. Thus, resetting back to GCT 0 was incomplete if hydrocortisone was applied after GCT 0, in the first half of the cycle, whilst full resetting was achieved if hydrocortisone was applied any time after GCT 10 (**Fig.6a**). This phase-dependent effect is known as a weak or ‘Type 1’ response and is reflective of the real-world ‘tug-of-war’ between cues that would occur with shift work or jet lag^37^. In affected individuals, for example, cortisol peaks may conflict with mistimed sleep, meal times, or light exposure. Critically, the impact of this temporal conflict was evident in the relative amplitude, damping, and robustness of organoid rhythms (**Fig.6b, Extended Data Fig.7a**); these parameters were optimised in the context of a ‘lie in’ (hydrocortisone at GCT 2.5), but worst with simulation of a ‘night shift’ (hydrocortisone at GCT 10). As a second proof of concept, we modelled clinical interventions in critical care, where synthetic glucocorticoids are often used at high doses^57^, and Targeted Temperature Management (TTM) practices may involve therapeutic hypothermia (e.g. 32°C) or enforced constant normothermia (37°C). We thus explored the effect of more natural (hydrocortisone) and synthetic (dexamethasone) glucocorticoids on bioluminescence rhythms, as well as MR and GR antagonists (S and X), in the context of temperature cycles followed by constant 32, 37, or 40°C (**Fig.6c, Extended Data Fig.7b**). Remarkably, we observed a near-complete loss of rhythm when organoids were cycled from 32 to constant 37°C in the presence of dexamethasone (**Fig.6c**). Moreover, dexamethasone and X, and hydrocortisone and S, had similar effects on rhythmicity after the shift to 32°C (**Fig.6d**), supporting the concept that the more natural glucocorticoid has some MR-mediated activity. Consistent with our earlier results (**Fig.3**), it appears that dexamethasone (commonly used in the clinic and by chronobiologists) is a poor substitute for cortisol with respect to brain tissue rhythms, and that modulation of GRs in isolation has a particularly detrimental effect on these rhythms under simulated therapeutic hypothermia. Using sucrose, we further modelled a clinically-applied hyperosmotic shift. At earlier stages of differentiation, sucrose improved robustness if applied at GCT 12, but had the opposite effect at D71, abrogating the robustness advantage achieved by hydrocortisone (**Fig.6e-f, Extended Data Fig.7c**). Finally, we confirmed the period-lengthening effect of the commonly prescribed antipsychotic lithium chloride (**Extended Data Fig.7d**). Overall, we have shown the utility of cerebral organoids to model clinically-relevant disruption and modulation of circadian rhythms in human brain tissues.

**Fig. 6.**
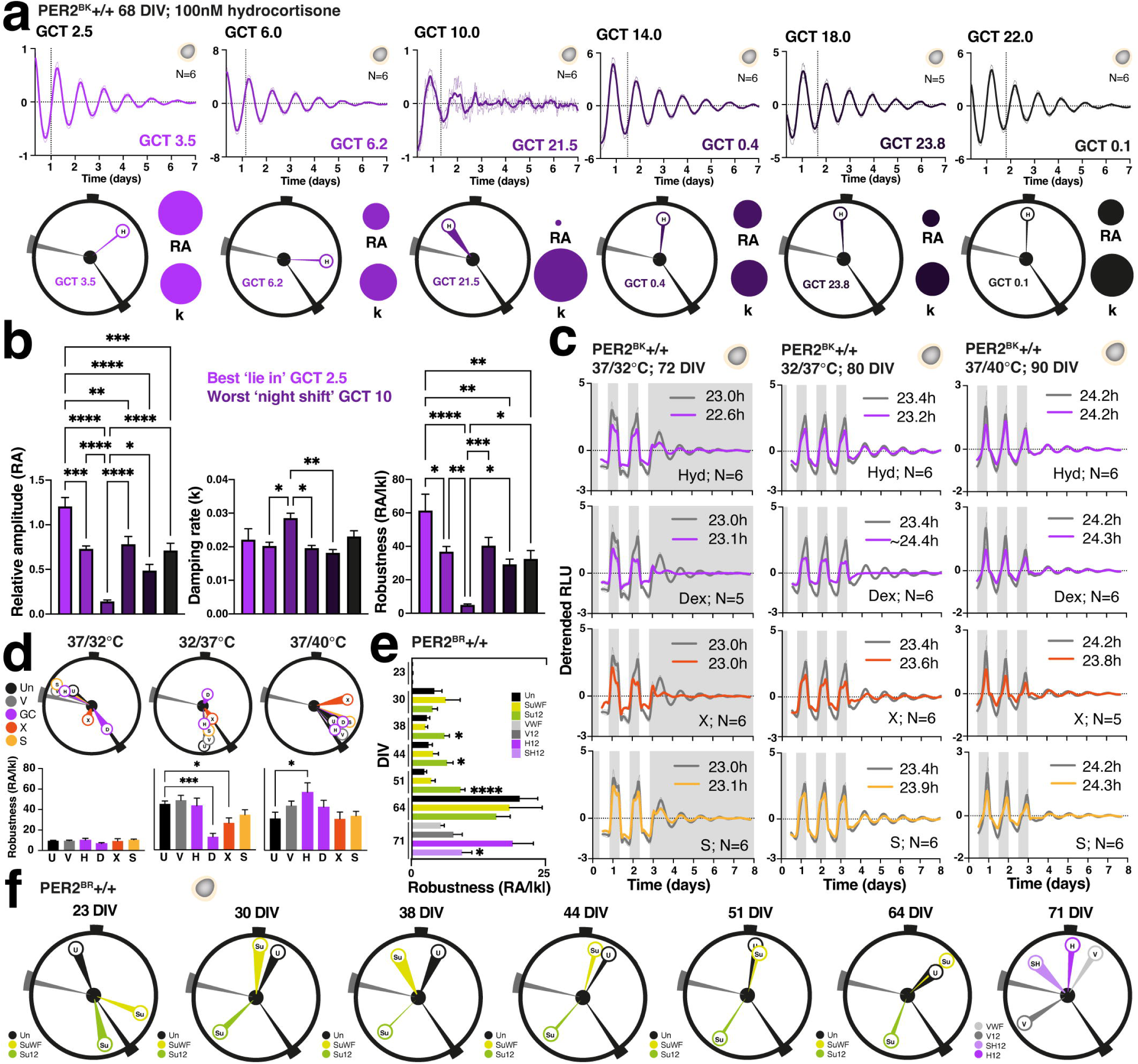
Modelling the disruption of human brain clocks. **(a)** Modelling ‘shift work’ with incorrect timing of glucocorticoid relative to underlying rhythm of cerebral organoids. After synchronization with media change, N=6 organoids were treated with hydrocortisone (H) at six different phases (GCTs in black) before bioluminescence was recorded under constant conditions at 37°C. Resultant phases are shown as colour-coded GCTs at bottom of each graph and mapped onto circadian clocks; size of adjacent bubbles depict relative amplitude (RA) and damping rate (k). Dotted vertical lines denote resultant phases, shifted along the x-axis by 4h with each successive treatment time. **(b)** Quantification of RA, k, and robustness from **(a)**. One-way ANOVA with Tukey’s correction for multiple comparisons, *p<0.05, **p<0.01, ***p<0.001, ****p<0.0001. **(c)** Organoid rhythm modulation by clinical interventions (temperature control and glucocorticoid). Effect of more natural (Hyd = 100nM hydrocortisone) and synthetic (Dex = 100nM dexamethasone) glucocorticoids and antagonists (X = 10uM exicorilant^48^, S = 10uM spironolactone) on rhythmicity in context of temperature cycles followed by constant 32, 37, or 40°C. White background denotes the higher temperature and vertical dotted lines show phase 24h after the last temperature transition. Period lengths are provided using fits of grouped data. An approximate period length is reported for the Dex condition in the middle panel; on manual inspection of individual traces, 2 of the organoids retained rhythmicity at low amplitude, 3 were just about discernible and 1 trace was better fit with a straight line. **(d)** Phase and robustness quantification of **(c)**. Note that control conditions set circadian phase to the predicted ‘shift to cold’ which is opposite to the ‘shift to hot’–most compounds tested have no impact on this, but dexamethasone (D) appears to invert the phase at 32°C. Both D and the GR-selective antagonist X significantly reduce the robustness of rhythms at 37°C, whilst hydrocortisone (H) increases robustness at 40°C. U or Un = untreated; V = vehicle (0.1% DMSO). One-way ANOVA with Tukey’s correction for multiple comparisons, *p<0.05, ***p<0.001. **(e)** Organoid rhythm modulation by hyperosmotic treatment. 100mOsm increase with sucrose improves robustness when given at GCT 12 (Su12) from 38-51 DIV. At 71 DIV, this same hyperosmolar intervention abrogates the robustness advantage achieved by hydrocortisone (Un = untreated, SuWF = sucrose with feed, VWF = vehicle with feed, V12 = vehicle at GCT 12, H12 = hydrocortisone at GCT 12, SH12 = sucrose and hydrocortisone at GCT 12). One-way ANOVA with Tukey’s correction for multiple comparisons, *p<0.05, ****p<0.0001. **(f)** Phase quantification of **(e)**. Sucrose alone has limited impact on phase after 44 DIV but partially abrogates the phase-shifting effect of hydrocortisone given at GCT 12 at D71. Note phasing is similar to that observed in Fig.3 for spironolactone applied prior to hydrocortisone. Compare data with **Supplementary Fig.6**.

## Discussion

Paramount to tackling the global burden of brain disease^58^ is a better understanding of how the brain works, and this demands a multi-dimensional approach. Through a combination of strategies, we have characterized circadian oscillations in 2D and 3D human neural model systems, thus demonstrating SCN-independent circadian rhythms in the neural cells of a diurnal mammal^31,59–60^. Several proteins that are linked to chronic brain disorders are highly dynamic within the cell; their spatial distribution and post-translational modification is temperature-sensitive across the range of brain temperatures observed in healthy humans^9,61^. By developing an array of clock reporter platforms, we have extended this dynamism to the fourth dimension of the daily clockwork, establishing that human neural physiology is intrinsically circadian and responsive to variations in temperature, glucocorticoids, and other timing cues that brain cells would receive *in vivo*^62^. It follows that the rhythms we observe under constant conditions are almost certainly enhanced by the cyclical brain environment of daily life. Whilst many diurnal variations in human physiology and behaviour may be paced by a principal central clock^63^, circadian oscillations in human brain cells can evidently exist without it.

Contrary to the traditional TTFL paradigm of the circadian molecular clockwork^15^, human cerebral organoid rhythms in PER2 translation prevail even when rhythms in the *Bmal1*:Luc transcriptional reporter are undetectable. Transcriptomics will help clarify whether this represents a true lack of rhythmic *Bmal1* expression or that the nature of the lentiviral construct renders the transgene unstable in human neural cultures. Importantly, the same *Bmal1*:Luc reporter faithfully illuminates robust circadian rhythmicity in other human cell types, including skin fibroblasts differentiated from the same NGN2 opti-ox™ stem cells that were used for organoid generation (**Supplementary Fig.1d**)^36–37,64^. Currently, these data suggest that circadian rhythms in *Bmal1* transcription may be poor and/or uncoupled from PER2 translational rhythms in human cerebral organoids.

Relative amplitude is a well-defined and measurable circadian parameter, but its biological relevance remains poorly understood. In a bioluminescent cell population, a reduction in rhythmic amplitude may represent a true loss of rhythmicity, or simply desynchrony of high-amplitude rhythms between individual cells^31^. Our current platforms do not reliably distinguish these possibilities, which requires single-cell microscopy at a much lower throughput. A further limitation is that our standard bioluminescence recording chamber controls only the incubated environment and not the immediate cell culture environment that cells and tissues are plated in. Refinement of this system with constant perfusion will enable further control of culture conditions, including more precise and physiological manipulation of timing cues and drug treatments^65^. Finally, since the relative proportion of dividing cells varies during organoid development, it is not yet possible to quantitatively dissect the impact of this variable on bioluminescent rhythmic parameters. Excepting paradigms that were development focused (**Fig.3**), circadian experiments were conducted at a stage when the vast majority of cells in each organoid are expected to be post-mitotic^66^. Nonetheless, we have charted the development of glucocorticoid resetting of circadian rhythms in cerebral organoids, capturing a time window during which this emerges and when it appears to ‘mature’. This parallels the relative expression of steroid receptor subtypes, intracellular steroid conversion enzymes, and drug efflux pumps described for the brain elsewhere^12–14^. Synchronization of organoid clocks with glucocorticoid emerged before convincing resetting to other timing cues such as temperature shifts, and hydrocortisone readily dominated phase and amplitude responses throughout differentiation (**Fig.3, Extended Data** Fig.4**, Supplementary Fig.3b**). Concordantly, we found that MR/GR inhibition later enhanced the resetting response to other timing cues in the media (presumably by removing any conflict from glucocorticoid, and enabling growth factors to dominate, **Supplementary Fig.3c**). Of importance to chronobiologists and clinicians, who routinely use dexamethasone for synchronizing circadian clocks and treating patients, respectively, we have highlighted the results obtained with different glucocorticoids during organoid development^12^. Selectively inhibiting GRs and MRs has produced data that suggest MRs play a role in neural circadian responses to cortisol, but not dexamethasone. Like the core TTFL components, most of the relevant targets for glucocorticoid sensing, conversion, and transport (NR3C1, NR3C2, HSD11B1/1L/2, ABCB1) were either not detected in our organoid mass spectrometry, or were not detected in sufficient samples to draw quantitative conclusions about temporal variation. An exception was ABCC1, which exports corticosterone (but not cortisol) from the brain^67^ and was not rhythmic in abundance. This is consistent with our ability to repeatedly synchronize organoids with hydrocortisone. The speed with which hydrocortisone effected resetting of PER2::LUC rhythms, and that this was completely blocked with antagonists applied just 30 min beforehand, concords with a non-genomic steroid activation of GRs and/or the phylogenetically older MRs^65,68–71^ that might help resolve the apparent decoupling of neural PER2 translation from the canonical TTFL model.

Strikingly, the phase-resolved relative amplitude of organoid proteins and phosphopeptides was reminiscent of a ‘cantilever’ distribution (**Fig.4g**), whereby comparative rigidity of the proteome at GCT 6 was complemented by greater variance in the phosphoproteome at this phase, whilst the opposite was true at GCT 18. This implies an intriguing dichotomy in which one half of the neural circadian cycle is dominated by changes in protein phosphorylation, whilst the other is dominated by variation in protein abundance. Irrespectively, we observed clear temporal consolidation of biological processes with prioritization of protein synthesis around the predicted temperature increase of the human ‘active phase’ (GCT 6) and neural signalling towards the end of this (GCT 15)^37^. Although we cannot directly map these phases to human behaviour, the quiescent part of the cycle in terms of functional enrichment (GCT 21) is predicted to align with mid-sleep^53^. A cellular basis for sleep remains controversial, and we caution against extrapolating our circadian findings to this highly complex neural process involving multiple brain structures^28,72^. We note only that the abundance and dynamism of protein phosphorylation in cerebral organoids is clearly biased towards one half of the circadian cycle, and that most of the synaptic phosphoproteins related to sleep need and sleep phenotypes are rhythmic in our dataset, including CaMKIIα/β at T286/T287 (**Extended Data Fig.5d**)^72–74^. RAIN outputs were prioritized in our analysis because manual exploration of the data showed that many rhythmic proteins and phosphopeptides had a distinctly non-sinusoidal and asymmetric pattern (**Extended Data Fig.5c**). We did however compare our results with those generated by other algorithms, finding that the proportion of rhythmic proteins varied from 13.1 to 38.6%, depending on the algorithm chosen, and using p-value cut-offs routinely applied in the field^75–77^. The 33% rhythmic proteome we report is higher than the 17.2% reported for the mouse forebrain^53^, whilst our 27% rhythmic phosphoproteome is broadly consistent with the 30% obtained from mouse forebrain synaptoneurosome^50^. Such similarity is surprising given the different species, temporal resolution, and bioinformatic approaches used, as well as markedly different paradigms (mice under entrained conditions for 24h versus cerebral organoids under constant conditions for 48h). Direct quantitative comparisons between these studies are therefore not appropriate. In the absence of a field-wide consensus on gold-standard methods for circadian neural mass spectrometry, we consider RAIN to best capture biologically- and neurologically-relevant waveforms that are challenging to fit with conventional cosinor modelling (**Fig.5**) and are likely to be missed with other tools^51–52^.

Beyond proteostasis, every aspect of brain physiology from cognition to pain perception and blood-brain barrier permeability varies by time of day^78–80^. Supporting the temporal enrichment of organoid proteomics, our MEA results indicate that the functional activity of human brain cells is inherently circadian^81^. It is perhaps unsurprising, therefore, that several neurological conditions exhibit time of day variation in symptom expression, including epileptic seizures, cluster headache, and migraine^82–83^. In the real world, the incidence of these episodic events will be further modulated by diurnal physiology that amplifies or entrains the neural clockwork e.g. daily brain temperature rhythms (**Fig.2**)^9,84^. Recordings from nocturnal rodents highlight that most extra-SCN brain regions display circadian rhythms in electrical activity that are in phase with locomotor activity (and higher body temperatures), but antiphasic with SCN electrical activity^30^. In human ALI-COs, the peak of spiking activity occurred at around GCT15, when the brain is expected to be cooling down, but peaks of bursting activity occurred in the early part of the circadian cycle, around the predicted wake time^37^. This phasing of burst activity was similar in organoids, where spiking peaks were, by contrast, well aligned with bursts. Further work is needed to decipher the physiological significance of these phase relationships and whether they correlate with sleep/wake patterns *in vivo*. Meanwhile, the phase-shifted circadian rhythms we describe in human astrocytes as well as the impact these cells have on neuronal rhythms and electrophysiology lends further weight to the notion that glia play an important role in neural timekeeping^85–86^. Importantly, however, the phase difference between human astrocytes and our other neural cultures is roughly equal (4-6h) but opposite (phase advanced) to those described in the mouse SCN (phase delayed)^85^. Notwithstanding differences in the choice of clock reporter, such a result may reflect an SCN-specific delay of astrocytic rhythms, or that the antiphasic relationship of nocturnal mouse versus diurnal human astrocytes contributes to the extreme contrast of behavioural chronotype between these two species. Exploring the resultant phase in SCN-specific versus cerebral mouse astrocytes co-cultured with or without human astrocytes would help clarify this.

Finally, we showed that the mistiming of clock resetting by hydrocortisone (as might happen in shift work, jet lag, or disease), readily disrupts the timing, amplitude, and robustness of human neural circadian rhythms. Of relevance to neurocritical care, further study should explore the interaction of glucocorticoid and temperature^9,87–88^ (**Fig.6c**) and how these cues are integrated by the clock, with the potential for ‘constructive’ and ‘destructive’ interference^37^. Whether circadian disruption plays a causal role in the development of chronic brain disorders remains to be formally tested, but changes in diurnal activity are linked to substance misuse and often predate episodes of mental illness^89–90^ as well as the onset of dementia-related symptoms^23^. Plausibly, a mismatch between external timing cues and the normal temporal order of intracellular processes (**Fig.4f**) presents a repetitive challenge to neural proteostasis that ultimately fails.

Target-led drug discovery in neurology, and particularly for neurodegeneration, has focused on correlates of disease typically modelled in a static fashion, based historically on definitive diagnoses that rely on a fixed (postmortem) endpoint. Our mass spectrometry data clearly show that many of these key targets are oscillating daily within human cerebral tissues; this variance should be considered at every stage of therapeutic development and during treatment regimens to optimise efficacy and safety, and reduce waste^91–92^. The tractable systems presented here can be deployed for high-throughput screening, but also to examine the impact of disease-causing mutations, diurnal physiology, and acute and chronic stressors^93–95^ on the workings of neural circadian clock.

## Methods

Methods are available in Supplementary material.

## Acknowledgements

This work was funded by the Medical Research Council, as part of United Kingdom Research and Innovation (MRC Clinician Scientist Fellowship MR/S022023/1 to N.M.R and MC_UP_1201/9 to M.A.L.). C.M.P. gratefully acknowledges funding from the Biotechnology and Biological Sciences Research Council David Phillips Fellowship (BB/T009314/1). N.K. acknowledges funding from the UK Engineering and Physical Sciences Research Council Centre for Doctoral Training in Sensor Technologies for a Healthy and Sustainable Future (EP/S023046/1). Cryopreservable wild-type iPSC-derived astrocyte progenitor cells were originally generated by Andrea Serio and expanded by N.M.R under the supervision of Siddharthan Chandran with support from Wellcome (096409/Z/11/Z). The opti-ox™ enabled microglia iPS cell line and reprogramming protocol were obtained from bit.bio Ltd. (Cambridge UK) under a research collaboration agreement. bit.bio Ltd. holds an exclusive license to opti-ox via Cambridge Enterprise. We thank Iván Shlamovitz and Ana Tufegdžić Vidaković for guidance on RNA sample processing, Catarina Franco of the LMB Mass Spectrometry Facility for guidance on protein sample preparation, and Joe Troughton (University of Cambridge) for assistance with the ALI-CO MEA cartoon. We thank the team at Cairn Research, Will Deacon at Axion Biosystems, Hazel Hunt at Corcept Therapeutics, members of the LMB Electronics and Mechanical workshops for technical support, and the UK Dementia Research Institute at the University of Cambridge for equipment support. Schematics were created using resources developed by the LMB VisLab. We thank John O’Neill for support and advice on circadian experimental design. We thank members of the Hastings Lab, and additional members of the Kotter Lab, Lancaster Lab, and O’Neill Lab for discussion and suggestions.

## Author information

### Author contributions

N.M.R. conceived the study, and C.P., M.K., and M.A.L. supervised the study. N.M.R. generated bioluminescent human stem cell lines with support from A.M. and A.D.B. for gene editing constructs, and support from M.A.S on nucleofection strategies. N.M.R. differentiated human neural cultures from transduced and edited stem cells. M.A.S. generated organoids from stem cells and validated their cerebral identity under the guidance of M.A.L. N.M.R. designed and performed all circadian experiments, with support from K.B. and N.P. for iAstrocyte validation and K.B. for Axion MEA work, and support from D.L-D.S and N.K. for ALI-CO and bespoke MEA work. N.M.R., K.B., N.K., and N.P. analysed data. S.Y.P-C processed the samples for mass spectrometry. A.Z. and E.S. assisted with proteomics and phosphoproteomics analysis. K.K. helped with validation of the iNeuron platform. N.K. engineered the bespoke ALI-CO MEA platform with the support of D.L-D.S under the supervision of C.P. and M.A.L. N.M.R. prepared figures and wrote the paper; M.A.S., K.B., D.L-D.S, and N.K. contributed images. All authors contributed to manuscript editing.

## Additional information

### Reprints and permissions information

For the purpose of open access, the MRC Laboratory of Molecular Biology has applied a CC BY public copyright licence to any author accepted manuscript version arising.

## Extended Data Figure Legends

**Extended Data Fig.1.**
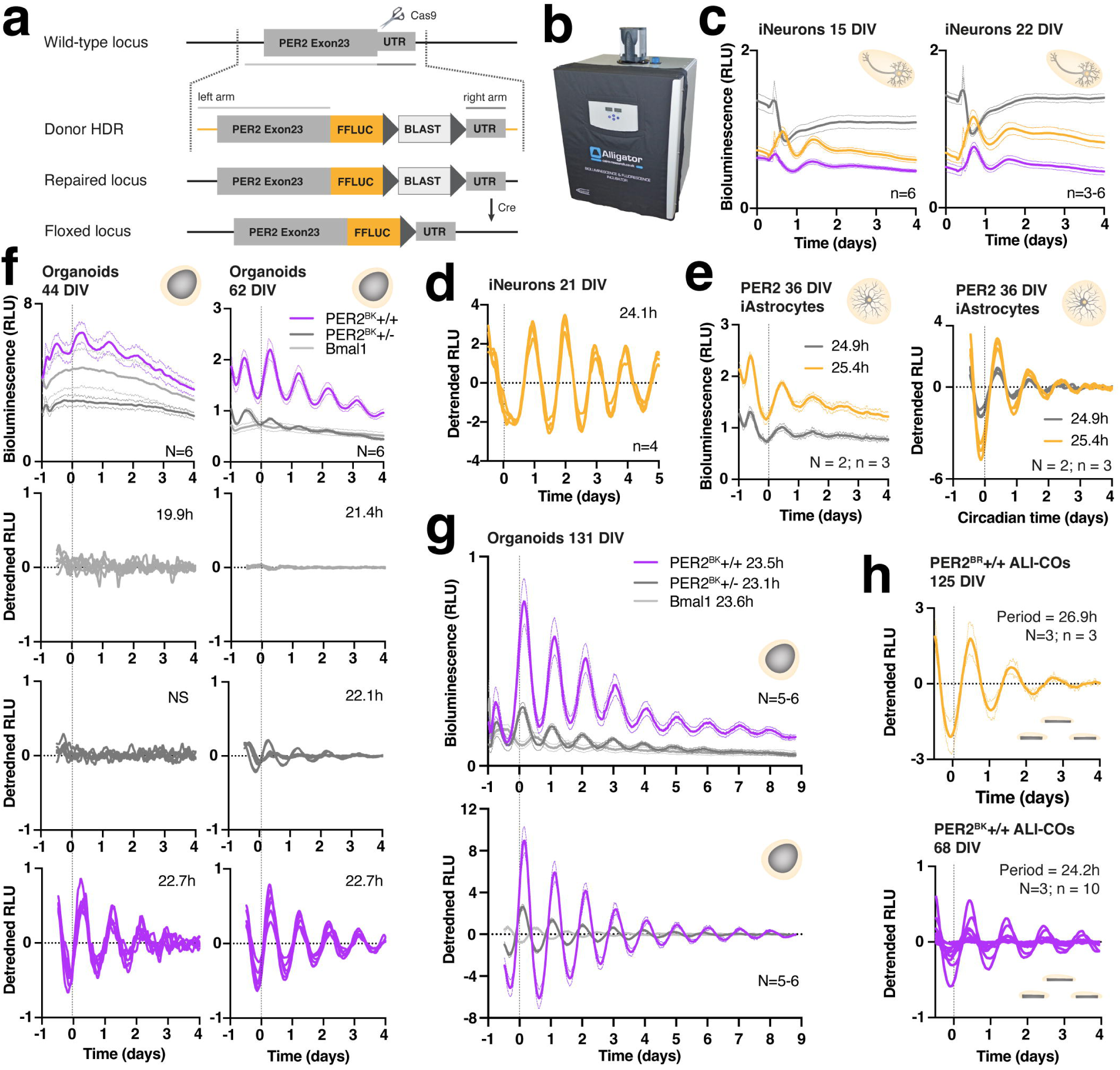
Related to Fig.1. **(a)** Schematic of human stem cell gene editing strategy to couple firefly luciferase (FFLUC) to PER2 translation with floxed blasticidin (BLAST) selection cassette. **(b)** Cairn Research ALLIGATOR used to collect bioluminescence data from human neural cultures. **(c)** Smoothed (non-detrended) bioluminescence data used to construct middle panel of Fig.1c. **(d)** Bioluminescence traces from individual wells of iNeurons used to construct lower right panel of Fig.1c; smoothed detrended data are shown. **(e)** Smoothed grouped data +/- SEM and detrended bioluminescence traces from individual wells of iAstrocytes used to construct Fig.1d. **(f)** Smoothed grouped data +/- SEM and individual organoid data to support Fig.1e. **(g)** Older organoids synchronized with 100% media change plus 100nM Hydrocortisone and upshift to constant 37°C for an extended 9 days. N=5-6 organoids per clone +/- SEM. Upper graph shows raw (background-subtracted), unsmoothed, grouped data; note similar phasing of bioluminescence peak in PER2 clones which contrasts with timing of peak in *Bmal1* bioluminescence. Baseline is substantially lower in monoallelic versus biallelic knockin PER2 clones, and even lower in *Bmal1* clone. Period lengths are shown. Lower graph shows 24-hour detrended and smoothed grouped data +/- SEM. Phasing of all clones at 0h is consistent with the combination of timing cues applied at −1 day (see later figures). **(h)** Additional data from older ALI-COs in upper panel sliced from PER2^BR^+/+ organoids (smoothed, detrended +/- SEM); lower panel shows data from individual slices used to construct Fig.1f.

**Extended Data Fig.2.**
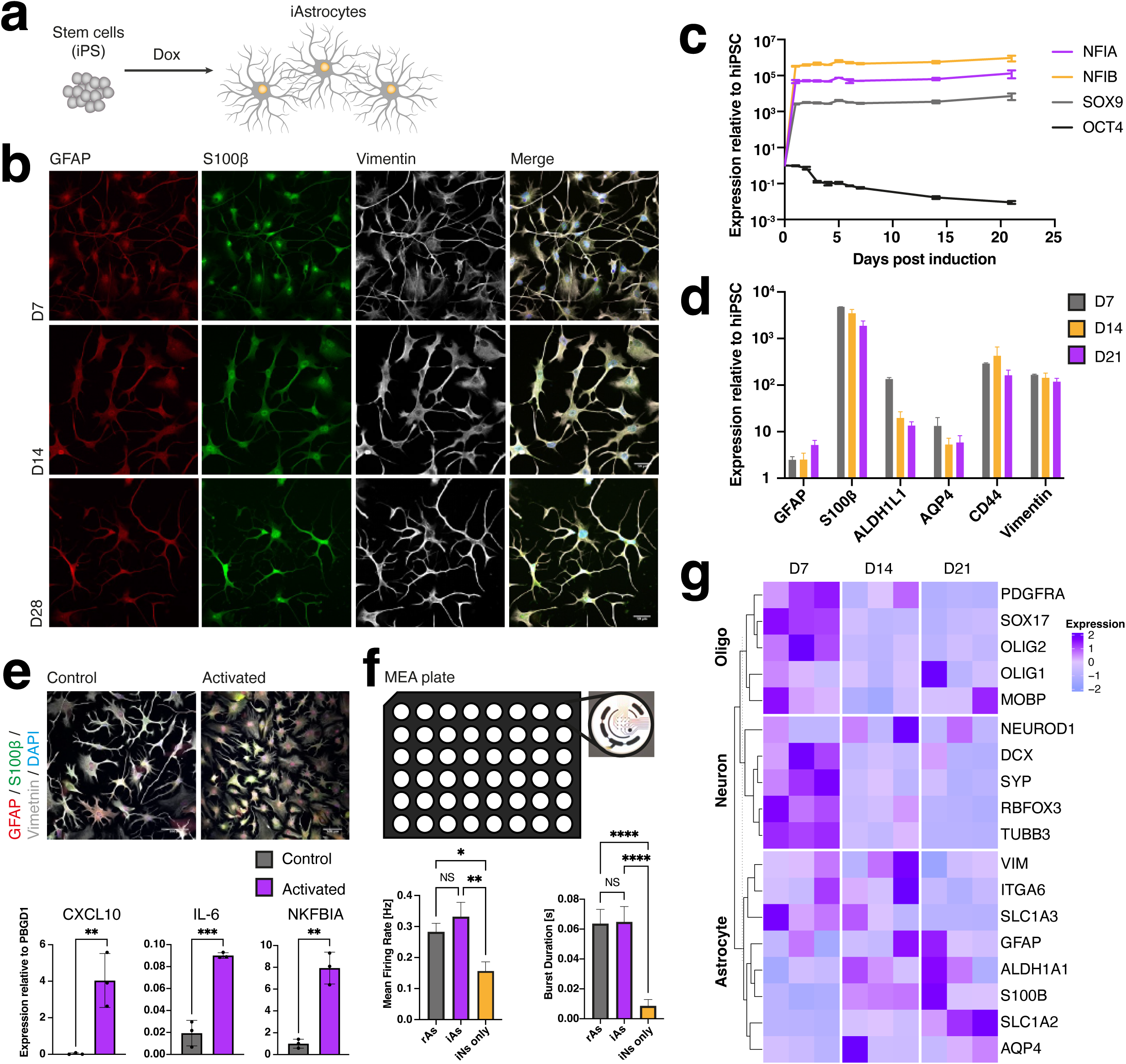
Related to Fig.1 **(a)** Experimental approach of generating iAstrocytes from opti-ox™ SOX9-NFIA-NFIB hiPSCs. **(b)** Immunocytochemistry staining for astrocytic markers GFAP (red), S100β (green) and vimentin (grey), at D7, 14 and 28 post induction. Scale bars: 50µm. **(c)** Quantitative RT-PCR analysis of mRNA levels for the exogenous transcription factors SOX9, NFIA and NFIB, and for the pluripotent gene OCT4. All data relative to PBGD1 and normalized to iPSCs. **(d)** Quantitative RT-PCR analysis of mRNA levels for the astrocytic markers GFAP, S100β, ALDH1L1, AQP4, CD44 and vimentin at D7, 14 and 21 post induction. All data relative to PBGD and normalized to iPSCs. **(e)** Cytokine activation assay based on IL-1β and TNFα treatment for 24h. Immunocytochemistry staining showing change in morphology from branched (control) to flat-like morphology (activated). mRNA levels of specific cytokines (IL-6, CXCL10 and NFKBIA) are increased following activation. Scale bars: 100µm. All data relative to PBGD1 and is presented as mean ± SD, student t test, **p<0.01, ***p<0.001. **(f)** MEA analysis of D21 iNeurons cultured with primary rat astrocytes or iAstrocytes and compared to control iNeuron monocultures. Data is presented as mean ± SD, One-way ANOVA, *p<0.05, **p<0.01, ****p<0.0001, NS=not significant. **(g)** Heatmap analysis of D7, 14 and 21 iAstrocytes based on genes enriched in neurons, oligodendrocytes, and astrocytes.

**Extended Data Fig.3.**
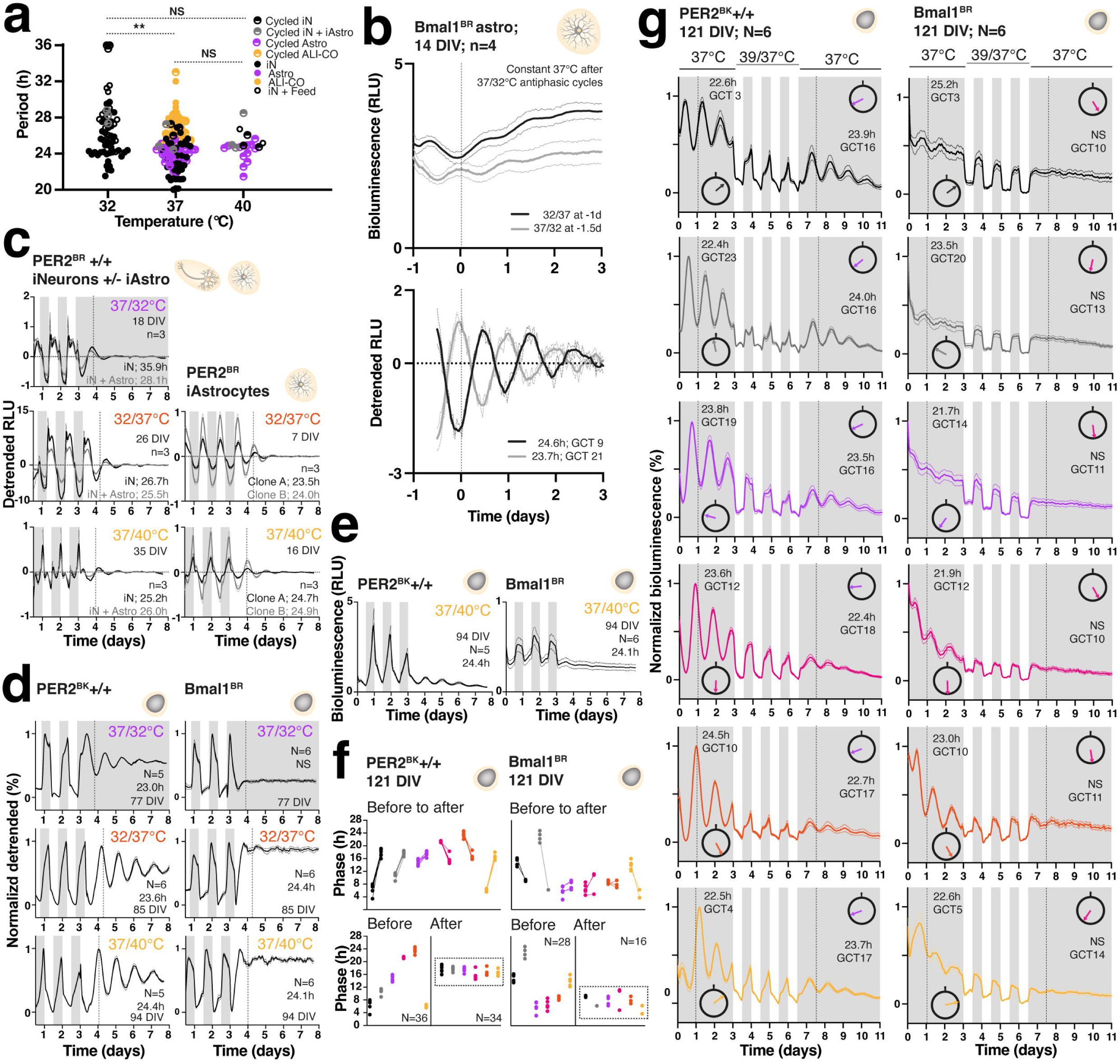
Related to Fig.2 **(a)** Temperature compensation of period length in other neural culture types to support Fig.2 **(a)**; the only significant difference was between 32°C and 37°C (**p= 0.004, Kruskal-Wallis with Dunn’s correction for multiple comparisons). **(b)** Opposing temperature entrainment of astrocytes followed by free run; vertical dotted line at time zero equates to 24h after start of constant conditions. Upper graph shows non-detrended data and legend denotes last temperature shift for each culture. Bottom graph shows same data detrended; legend denotes period lengths and phases (at time zero). Note antiphasic rhythms and that temperature drop sets phase to GCT 21 whilst temperature increase sets phase to GCT 9, both of which are around 3h phase-advanced compared to other cell types. 0.5mM luciferin was used with 12-h detrending and no amplitude normalization. **(c)** Temperature resetting data traces from iNeurons cultured with and without iAstrocytes used to construct wheels in Fig.2b. Note that fit for iNeurons alone after switch to constant 32°C was preferred over a straight line but is at the upper end of the fit constraint; result should be interpreted with caution. Period lengths represent an average of fit values for individual wells of cells. **(d)** Temperature resetting data traces from organoids used to construct Fig.2b; detrended data have been amplitude-normalized to aid visualization. Note that bioluminescence rhythms are more robust at higher (more physiological) temperatures and *Bmal1*:Luc rhythms are absent at 32°C. **(e)** Raw smoothed grouped data from the last temperature transition above without amplitude normalization; note differing scale of y-axis for organoids with different clock reporters (necessary for visualization). **(f)** Linear scale presentation of raw phase data points (prior to GCT conversion) from individual organoids used to construct Fig.2e. **(g)** Amplitude-normalized raw smoothed grouped data to support Fig.2d. Mini clocks highlight the phase of the rhythm (in circadian hours, thus normalizing for variable period lengths) before and after the temperature cycle, to the nearest circadian hour, relative to GCT 0 (black notch). Period lengths are displayed in hours where the cosinor fit was significantly preferred over a straight line for grouped data (otherwise NS) and mean GCT values of the resultant phase are given to the nearest circadian hour, whilst the x-axis displays external time. Vertical dotted lines signify phase of rhythm 24h after the start of constant conditions. For Bmal1 organoids, the quantification of phase (and thus GCT) included only individual organoids whose bioluminescence was significantly rhythmic. Colour coding corresponds to different phasing before the temperature cycle. Normalization of detrended data refers to normalization for amplitude during each stage of the experiment (before, during, and after temperature cycling).

**Extended Data Fig.4.**
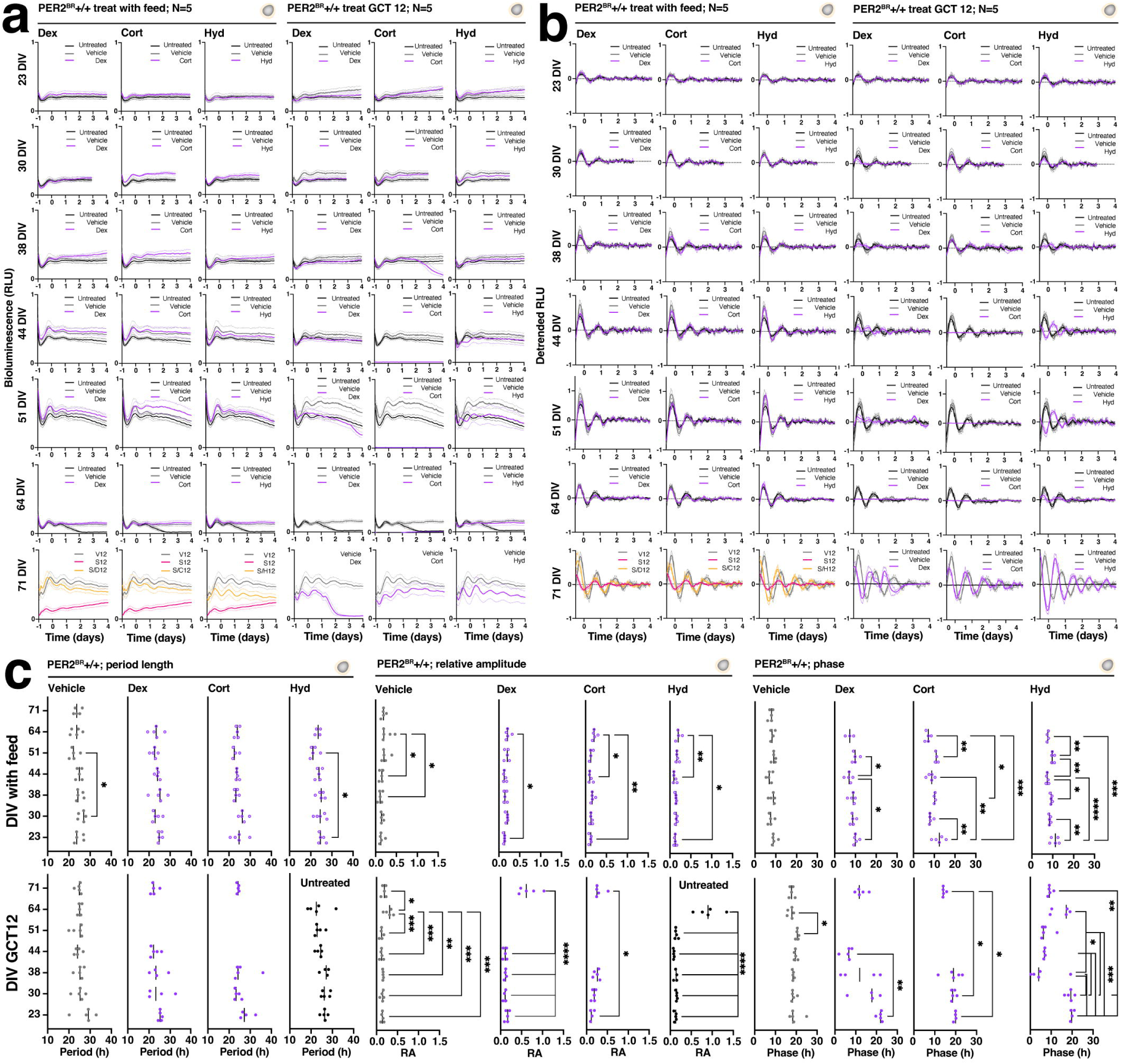
Related to Fig.3. **(a)** Raw smoothed grouped data for all treatments at each age in culture; acute drops in the baseline reflect a well of organoids (and thus a treatment condition) lost to fungal contamination at that time point and thereafter. For the final run at 71 DIV, the range of treatment conditions was condensed based on available wells remaining. Note progressive increase in baseline in the S-only condition. **(b)** Detrended versions of data in **(a)**. **(c)** Period, relative amplitude, and raw phase data (not converted to GCT) for individual organoids treated with different glucocorticoids at each developmental stage. The GCT 12 hydrocortisone period and relative amplitude data are already shown in Fig.3b and are replaced here with data from untreated control organoids. See **Supplementary Fig.3a** for ‘with feed’ phase data for untreated control organoids. One-way ANOVA with Tukey’s correction for multiple comparisons, *p<0.05, **p<0.01, ***p<0.001, p<0.0001.

**Extended Data Fig.5.**
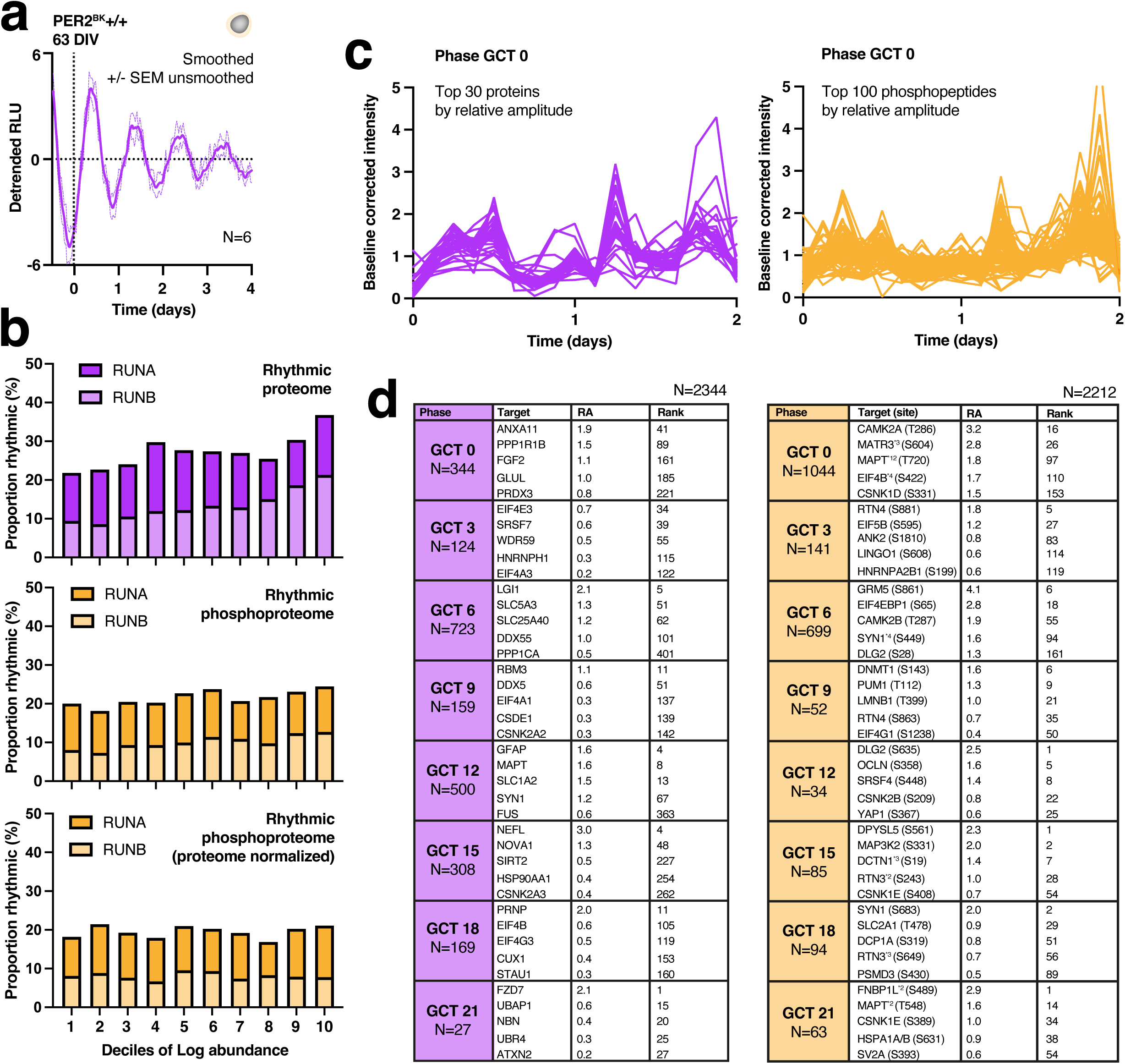
Related to Fig.4. **(a)** Smoothed detrended data to support Fig.4a. **(b)** Proportions of rhythmic proteins and phosphopeptides plotted by deciles of abundance; proportions for both batches (A and B) are shown on each graph. The middle graph plots the phosphopeptide data before normalization to protein abundance (lower graph). **(c)** Intensity data across the time course (corrected to baseline media intensity) for top 30 and top 100 hits for the proteome and phosphoproteome, respectively, in the GCT 0 cluster. **(d)** Selected brain disease, sleep, and circadian-relevant hits in the rhythmic proteome (left) and phosphoproteome (right) by phase of peak abundance together with relative amplitude and ranking within each phase. Numbers of hits per phase are shown in left column of each table and rank value identifies the position of that hit within the phase-clustered list. Asterisks highlight proteins that were rhythmically phosphorylated multiple times and/or at more than one site (superscript value denotes total number of rhythmic phosphopeptides in clustered list). For these asterisked targets, the RA and rank value is given for the highest amplitude phosphopeptide, which is specified by phosphosite in parentheses. Other notable hits (not shown) include EIF3D, phospho-RIMS1, phospho-ANK3, and phospho-CRTC1 at GCT 6, EIF2A, phospho-ANK3 at GCT 9, SLC1A4, DNM1, DNM3, PPP2R5B, PPP2R5E, PPP3CB, RTN3, RTN4, GSK3B, HSPA5 at GCT12, RIMS1, ANK3, and GRM5 at GCT 15, CRTC1 at GCT 18, and phospho-DDX3X, phospho-ANK3, and phospho-LRRC7 at GCT 0^72–74,98–99^. The sleep-promoting kinase SIK3^100^ was detected in our proteomic and phosphoproteomic datasets but did not meet the cut-off for rhythmicity in either according to RAIN^51^.

**Extended Data Fig.6.**
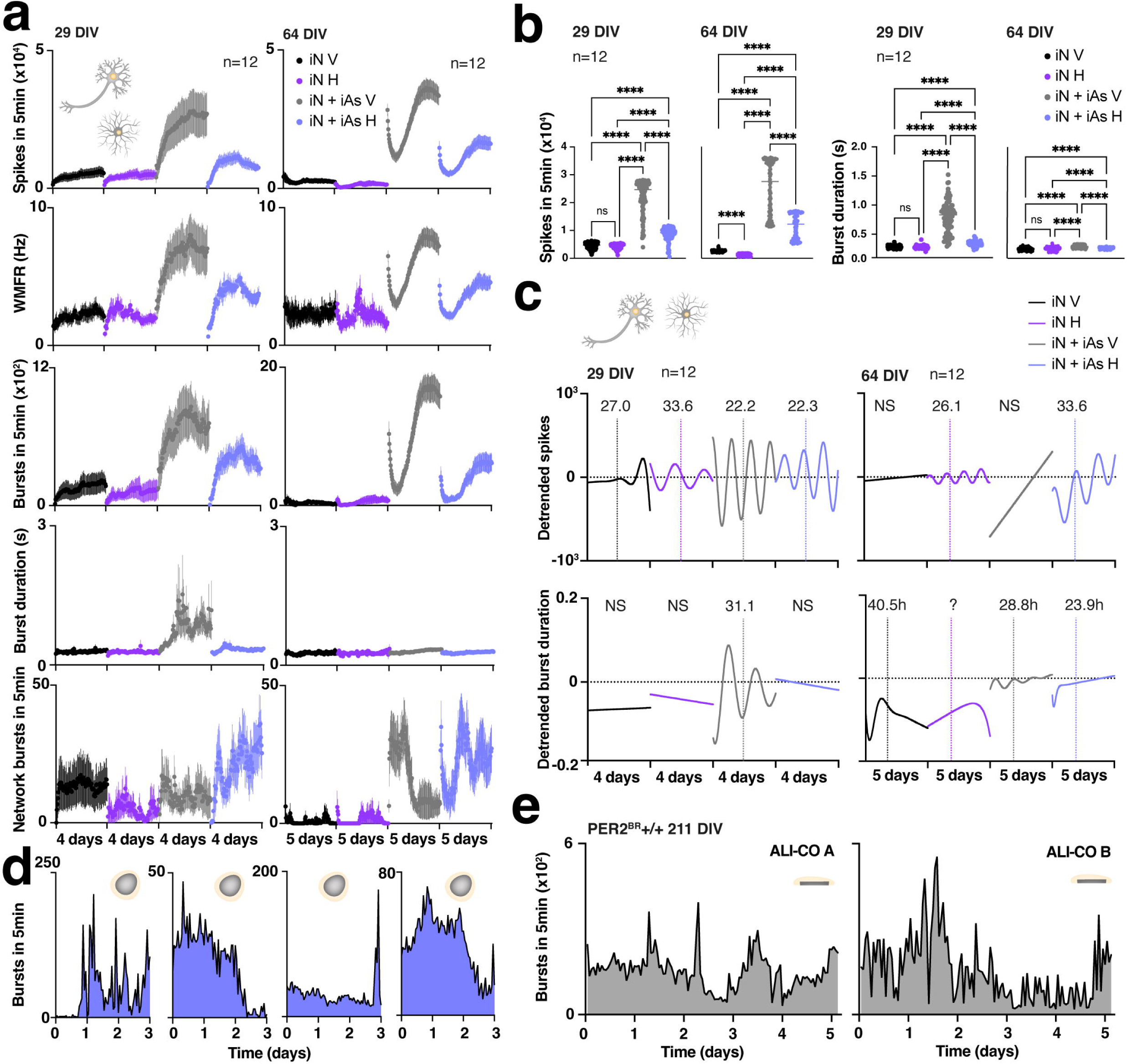
Related to Fig.5 **(a)** Raw Axion MEA data from iNeuron and iNeuron-iAstrocyte co-cultures (without smoothing or detrending) to support Fig.5c. **(b)** Summary spike and burst duration data to support Fig.5b. **(c)** Nonlinear fits of detrended spike and burst duration data. **(d)** Raw Axion whole organoid burst data to support Fig.5d. **(e)** Raw ALI-CO burst data to support Fig.5e.

**Extended Data Fig.7.**
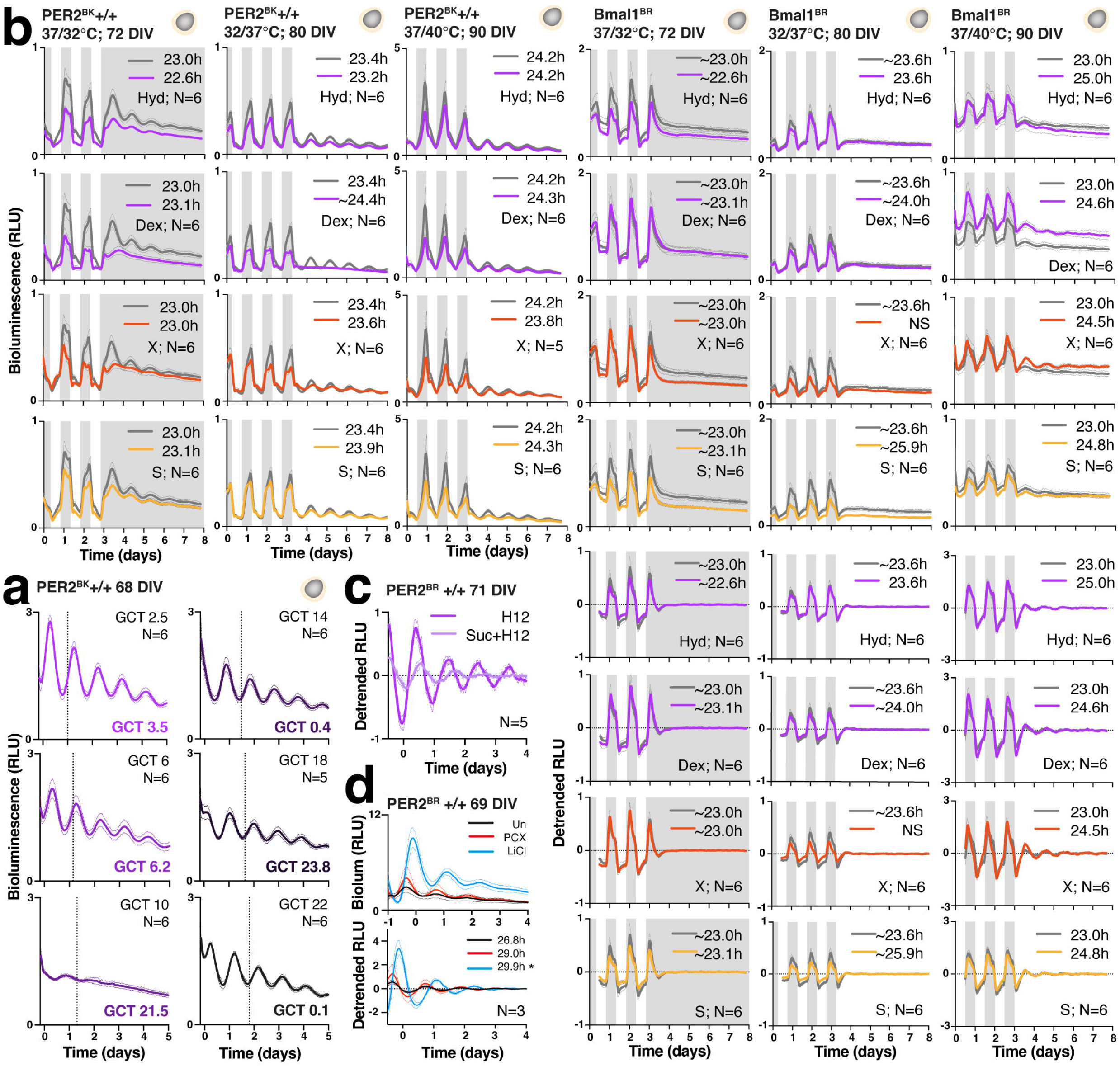
Related to Fig.6. **(a)** Raw smoothed grouped data +/- SEM used to construct Fig.6a. **(b)** Raw smoothed grouped data +/- SEM to support Fig.6c plus data from *Bmal1*:Luc organoids collected in parallel; note that *Bmal1* rhythms are generally poor but best observed at 40°C. Period lengths are presented for the fit on grouped data and do not imply that each individual organoid in that group/condition was significantly rhythmic. Where a straight line was preferred for two or more organoids in that group a tilde (∼) is denoted for the period length and the absolute value should be interpreted with caution. NS = not significant for grouped fit. **(c)** Smoothed detrended data +/- SEM of unsmoothed data to support result in Fig.6e. **(d)** Raw smoothed and detrended data showing higher baseline and period lengthening effect of 30mM LiCl versus untreated organoids (*p=0.03, unpaired t test with Welch’s correction). Red traces refer to rhythms in organoids after washout from treatment with 300uM picrotoxin. Un = untreated, PCX = picrotoxin.

## Supplementary information

**Supplementary information** The online version contains supplementary material available at XXX

**Supplementary Table 1.**
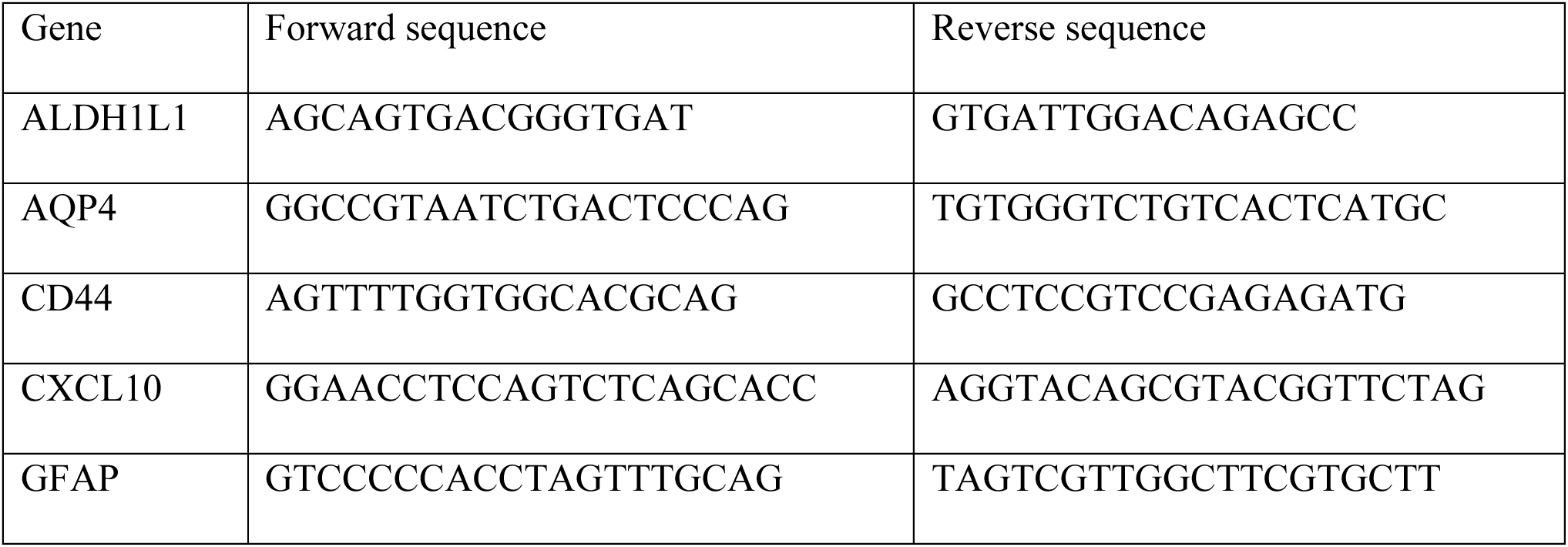
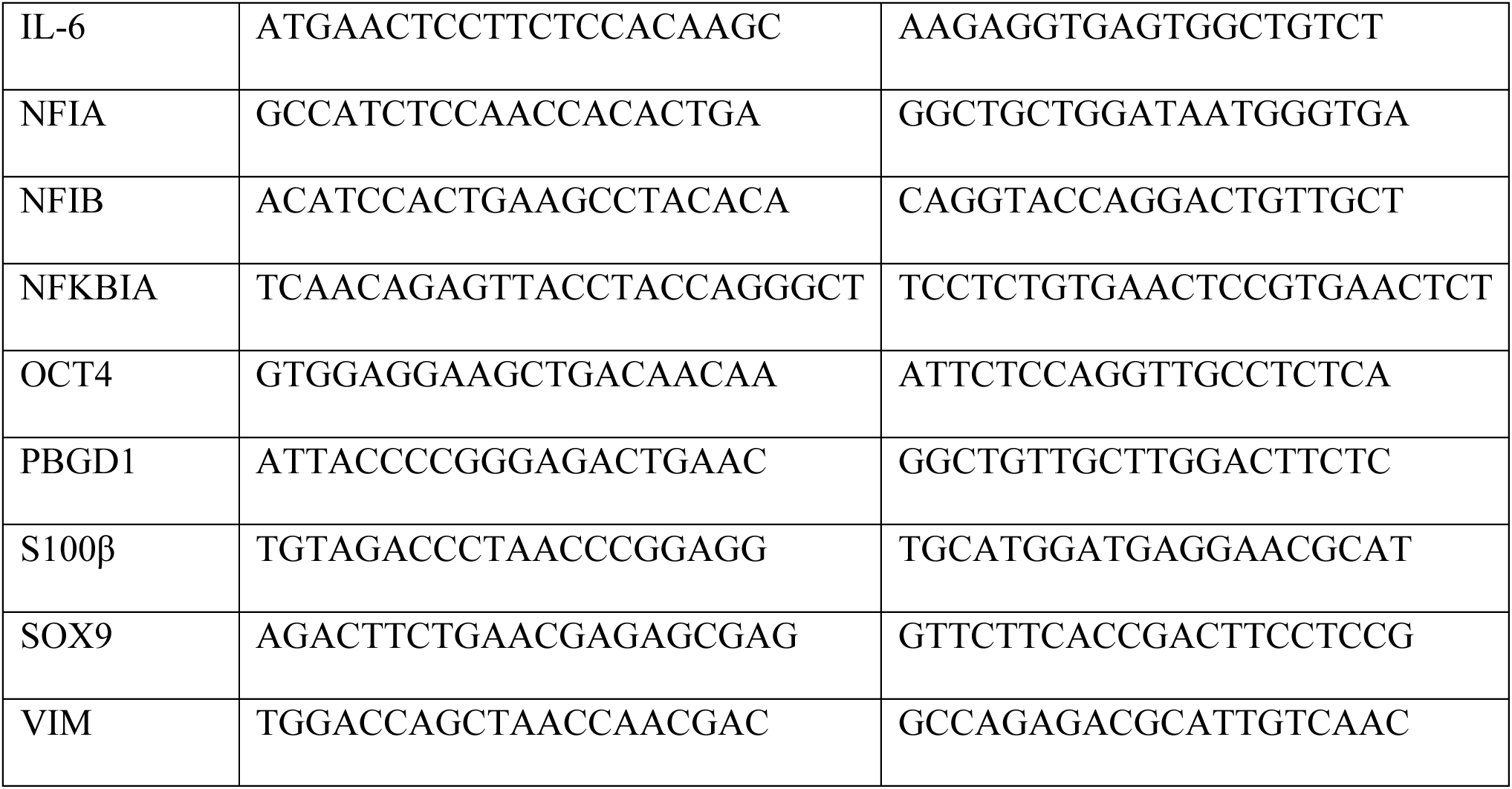
List of primers used for RT-qPCR analysis.

## Supplementary figure legends

**Supplementary Fig.1** Related to Fig.1 **(a)** Validation of CRISPR/Cas9 editing of endogenous PER2 locus by PCR. Left hand gel shows PCR products generated using 5’external primer set which flanks a region upstream of PER2 exon 23 and the region fusing exon 23 to firefly luciferase. The 1625bp product confirms successful edit in BR and BK clones with no product for WT. Right hand gel shows PCR products generated with long HDR primer set which flanks the start of PER2 exon 23 and 3’UTR region downstream of luciferase. The WT product appears at 203bp and is absent from BR and BK clones confirming absence of WT allele and successful biallelic knockin (with repaired product at 1904bp that is observed only after the blasticidin resistance cassette is removed). WT = wild-type non-bioluminescent NGN2 opti-ox™ stem cells, BR = PER2+/+ blasticidin-resistant, bioluminescent NGN2 opti-ox™ stem cells, BK = PER2+/+ blasticidin-knockout, bioluminescent NGN2 opti-ox™ stem cells. **(b)** Principal component analysis (PCA) of transcriptomic data demonstrating clustering of the iAstrocytes separate from hiPSC (D0). The percentage of variance explained by the PCAs is indicated between parentheses. **(c)** IHC validation of PER2^BR^+/+ cerebral organoids showing primarily dorsal forebrain identity with some choroid plexus; left-hand images from organoids at D73, right-hand images from organoids at D61 (TBR2 = dorsal forebrain intermediate progenitor marker, HuCD = neuronal marker, SOX2 = radial glia/progenitor cell marker, DLX2 = ventral forebrain specific intermediate progenitor marker, TTR = choroid plexus marker). **(d)** Smoothed detrended ALLIGATOR data recorded at 37°C from human skin fibroblasts differentiated from human Bmal1^BR^ bioluminescent NGN2 opti-ox™ stem cells. Dox = treated with doxycycline; n=3 wells of fibroblasts per condition. Note clear rhythms of appropriate period length here but relatively poor rhythms when these same stem cells were used to make iNeurons and organoids (Fig.1). The phase was set by full media change containing insulin and other growth factors at −1 day, with constant conditions starting at zero days. This phase (trough of *Bmal1*:Luc = GCT 6.9) is consistent with that set by growth factors^37^. Dox failed to transdifferentiate fibroblasts into iNeurons and had negligible impact on circadian rhythms.

**Supplementary Fig.2** Related to Fig.2 **(a)** Brightfield images of iNeuron monocultures cultured for 3 days at constant temperatures shown. Note healthy, viable cells at all temperatures. Scale bar 400um. **(b)** Raw (smoothed) and detrended data (without amplitude normalization) to support Extended Data Fig.3d.

**Supplementary Fig.3** Related to Fig.3 **(a)** Raw phase data (not converted to GCT) for ‘with feed’ untreated control organoids at each developmental stage. Phase reflects that set by full media replacement alone and by D64 corresponds to ∼GCT 2 (as predicted from organoid media supplements). At earlier stages, the phase sits closer to GCT 0, suggesting that phase setting by glucocorticoid emerges before other cues like insulin. One-way ANOVA with Tukey’s correction for multiple comparisons, *p<0.05. **(b)** Comparison of cue strength in young (D46) organoids which were cultured at constant 37°C or in 37/32°C temperature cycles before final shift to constant 32°C alongside treatment with vehicle (V) or 100nM hydrocortisone (H) without a media change. Recording was at 32°C. Note that prior temperature cycling has had negligible impact on relative amplitude, whilst H resets cultures to ∼GCT 23, irrespective of prior temperature conditions. This resultant phase reflects conflict between temperature downshift and glucocorticoid, with the latter being the stronger cue since cooling has delayed the phase by no more than 1h. Prior temperature cycling slightly suppressed the amplitude increase achieved with H. At this age in culture, temperature shift alone is a weak timing cue and fails to set organoids to the predicted ‘cold phase’ from Fig.2b data. **(c)** Washout at D95 after organoids treated at GCT 12 with hydrocortisone (H) or selective inhibitors (X or S) 30 min prior to hydrocortisone. The washout involved a full media replacement at 37°C, with recording at 37°C. Note that all rhythms are set to the same phase (∼GCT 8), consistent with resetting by growth factors such as insulin. Relative amplitude is greater in the context of prior GR/MR inhibition suggesting that this has heightened the response to other timing cues in organoid culture media. In **(c-d)** vertical dotted line at zero days represents phase 24h after start of constant conditions; the axis is displayed in standard time. Data are presented as smoothed and detrended +/- SEM of unsmoothed. RLU = relative luminescence units.

**Supplementary Fig.4** Related to Fig.4. Stages of early data processing for proteomics **(a)** and phosphoproteomics **(b)** including steps in chronological order from left to right. ‘Raw’ data include targets that were detected in all samples (all time points in both batches), ‘Corrected’ data correct for differences in submitted protein quantity prior to sample loading normalization (SL) and internal reference scaling (IRS) using the pooled sample data. In each case, data for batch A are shown in greyscale, and those for batch B in colour (purple = proteome, orange = phosphoproteome). Cluster plots, density plots, and boxplots are provided. In later steps, data for batch B are superimposed on those for batch A for density plots. The extra step for phosphoproteomics includes normalization to respective protein intensity. **(c)** Proportions of phosphorylated amino acids in the ‘all sample’ filtered (total) phosphoproteome (upper graph) and rhythmic phosphoproteome (lower graph). pS = phosphoserine, pT = phosphothreonine, pY = phosphotyrosine^50^.

**Supplementary Fig.5** Related to Fig.5 **(a)** Paradigm for terminal recording of 48-well MEA plate used for Fig.5. 10-minute recordings were performed with vehicle (DMSO) at baseline prior to washout and application of first compound with full media replacement, washout, application of second compound with full media replacement, washout, and finally TTX (a potent voltage-gated sodium channel blocker) with full media replacement. The plate was stabilized for 10 min at constant conditions (5% CO_2_, 37°C, humidified) prior to each recording under the same conditions, n=6 wells per condition for each culture type. Washes performed with Neurobasal media, whilst full media replacements included standard culture media for iNeurons, or 50/50 mix of standard culture media for iNeurons and iAstrocytes. **(b)** Summaries of raw data showing greater spiking and bursting activity in the presence of astrocytes, with negligible effect on burst duration. **(c)** Same data but normalized to baseline recording for each condition for better visualization. Note elevation of burst activity in iNeurons after CNQX treatment that remains elevated with picrotoxin, an increase in spiking and burst activity with bicuculline, loss of bursting after a toxic dose of glutamate, reduced firing frequency in the presence of GABA and rebound increase in firing with LiCl. In the presence of astrocytes, picrotoxin increases bursting activity, whilst the effects of glutamate are tempered, but bicuculline markedly increases burst frequency; GABA remains inhibitory and LiCl reverses this. Note complete lack of activity in all conditions after TTX treatment. B = baseline (vehicle) or bicuculline, V = vehicle (DMSO), C = CNQX, P = picrotoxin, G = GABA or glutamate, L = LiCl.

**Supplementary Fig.6** Related to Fig.6. **(a)** The same organoids used in **Supplementary Fig.3b** underwent a full media replacement and were shifted from 32°C to constant 37°C at D51. The organoids previously treated with hydrocortisone also received a hyperosmotic stimulus (+100mOsm sucrose) with the media change. Note that relative amplitude more than doubled for all conditions (compare y-axis with **Supplementary Fig.3b**) but a hangover effect remains for the original 32/37°C temperature-cycled organoids, whose amplitude does not reach that of organoids originally maintained at constant 37°C and the phase is slightly shifted. The hyperosmotic stimulus appears to have increased amplitude further, but prior hydrocortisone treatment may have influenced this. All rhythms are in the same phase initially, consistent with GCT expected for a temperature increase (6.5) or growth factor stimulus (7). Data are smoothed and detrended +/- SEM. Note x-axis is plotted here in circadian time to correct for differences in period length between conditions. **(b)** Raw grouped bioluminescence (upper) and smoothed detrended (lower) +/- SEM data from PER2^BR^+/- organoids fed at 37°C on D88 with or without a hyperosmotic stimulus (+100mOsm sucrose); note apparent phase shift but lack of amplitude enhancement. By contrast, the same hyperosmotic stimulus did not cause a phase shift in PER2^BR^+/+ organoids when applied with feed at D51 (see **Fig.6f**). F = feed, Su = +100mOsm sucrose.

## Supplementary videos

Supplementary videos will be available with the final version of the manuscript and can be viewed in the interim by contacting the corresponding author.

**Supplementary Video 1** Differentiation of iNeurons over 14 days captured with Incucyte® (Sartorius).

**Supplementary Video 2** Sped up video of bioluminescent PER2^BK^+/+ cerebral organoids (N=18) captured in ALLIGATOR. Pulsing circadian rhythmicity is immediate after synchronization with hydrocortisone; note damping of bioluminescence with loss of synchrony within and between organoids over time.

