## Supplementary material for "Circadian clocks in human cerebral organoids": Methods

*Cell culture and cell lines*

All stem cells and differentiated mono- and co-cultures, as well as 3D organoids and ALI-COs, were cultured in humidified conditions at 37°C and 5% $CO_2$ unless specifically stated otherwise. Neural culture media recipes and supplements are listed at the end of the methods section.

*Human pluripotent stem cells (hPSCs)*

Human embryonic and induced pluripotent stem cells (hESCs and hiPSCs, respectively)[1-3] were cultured on Geltrex™ (Thermo Fisher Scientific) or Matrigel® Basement Membrane Matrix (Corning, 356234) coated culture dishes in Gibco™ StemFlex™ Medium (Life Technologies; Thermo Fisher Scientific, A3349401) with 1% penicillin-streptomycin. Cells were passaged in small clumps using 0.1M EDTA or ReLeSR™ (StemCell Technologies) every 6-7 days at 85% confluence. RevitaCell™ (Thermo Fisher Scientific) was added to the culture media at 1X during procedures that induce cell stress (thawing, nucleofection etc). H9 (human female) embryonic stem cells purchased from WiCell (WA09) were approved for use by the UK Stem Cell Bank Steering Committee and were previously modified to generate NGN2 OPTi-OX hESCs as described[1]. A1ATD-hiPSCs were used for generating the original OPTi-OX iAstrocyte line[2-3]. Information relating to the hiPSC line from which the ioMicroglia™ model was generated is available with the product information on the bit.bio website. The line was created from skin fibroblasts (the donor was a caucasian adult male with normal karyotype 46,XY). hPSCs were cryopreserved in STEM-CELL BANKER® DMSO-FREE (AMSBIO) according to the manufacturer's instructions.

*Gene targeting of human iPSCs to generate iAstrocytes*

An optimized version of iPSC-derived iAstrocytes was generated based on a previously established OPTi-OX iAstrocyte line[3]. The expression cassette comprising of the 3 transcription factors, SOX9, NFIA and NFIB (S9AB), was cloned into the pAAV_HS4 vector (courtesy of Professor Cédric Ghevaert, Department of Haematology, University of Cambridge, UK). The pR26_CAG-rtTA and the pAAVS1_SOX9-NFIA-NFIB vectors were targeted into the human ROSA26 and the AAVS1 loci, respectively, by nucleofection as described previously[3–5]. Briefly, human iPSCs were dissociated into single cells with Accutase (Thermo Fisher Scientific), and $2x10^6$ cells were nucleofected using the P3 Primary Cell 4D-Nucleofector kit (Lonza) and cycle CA-137 of the Lonza 4D-Nucleofector System. Nucleofected S9AB-hiPSCs were plated onto Geltrex-coated dishes and cultured in StemFlex medium. Clone-R (StemCell Technologies) was added for 48h after nucleofection to promote cell survival. Drug-resistant S9AB-hiPSC clones from targeting experiments were screened by genomic PCR to verify site-specific transgene integration, to determine the number of targeted alleles, and to exclude off-target integrations (data not shown). Clones that were homozygous for both the rtTA vector and the reprogramming cassette were chosen for further expansion and subsequent experiments.

*Bmal1:Luc transcriptional clock reporter hPSCs, astrocyte progenitors, and skin fibroblasts*

For the production of lentivirus, the following were added to a sterile Eppendorf: X ul sterile $H_2O$ to 700uL final volume, 70ul 1.5M NaCl (0.22uM filtered), plasmid DNA (to a total of 7ug ), and 35ul PEI, added dropwise, vortexing between drops. Plasmids included 3.5ug transfer (pLV6-Bmal-luc https://www.addgene.org/68833/), 2.45ug packaging (psPax2 https://www.addgene.org/12260/) and 1.05ug envelope (pMD2.G https://www.addgene.org/12259/). The mixture was incubated at room temperature for 10 min then added dropwise to a 10cm dish of HEK 293T cells (seeded 2-24h previously, at approximately 60% confluence), tipping the dish back and forth to mix. The dish was incubated at 37°C for 48h and virus collected by removing and saving the media. Cell culture media was replaced and cells harvested again after 72 or 96 h if required. The virus-containing media was centrifuged at 10,000 rpm for 1 min and the supernatant aliquoted in 1ml aliquots and frozen at -70°C. Cells to be transduced were cultured to 50-80% confluence for the day of transfection. One control well was included as the non-virus transduction and selection control. Transduction media was prepared by diluting virus 1:10 in the regular culture media for the cell type and supplemented with 8ug/ml polybrene. All media was removed from each well and replaced with transduction media. Cells were incubated for 48 h at 37°C and the media was then refreshed with normal media. After 24 h, the media was replaced with selection media (5ug/ml blasticidin). Cells were checked daily for cell death. Once all cells had died in the control well but had begun to proliferate in transduced wells, the selection antibiotic was reduced to 2.5ug/ml thereafter to retain expression of the transgene during expansion and passage. Blasticidin was removed only during experiments. Wild-type iPSC-derived astrocyte progenitors were generated by Andrea Serio as described previously[6]. Skin fibroblasts were differentiated from *Bmal1*:Luc NGN2 OPTi-OX hESCs using a previously validated protocol[7].

*Gene targeting to generate human PER2::LUC clock reporter stem cells*

hPSCs endogenously expressing the PER2-LUCIFERASE were generated using CRISPR/Cas9 as described[8], except that a floxed blasticidin selection cassette was included in the HDR template (**Extended Data Fig.1a**). Briefly, the guide RNAs for Cas9 editing at the stop codon of PER2 were designed using a combination of CHOPCHOP v2[9] and ATUM gRNA design tool (atum.bio/eCommerce/cas9/input) and default parameters. Guide RNA sequences, lacking the PAM sequence were inserted as annealed complementary single-stranded oligos into BbsI-digested pSpCas9-2AGFP (PX458, Addgene). HDR templates, inserting the coding region lacking the initial methionine codon of human codon-optimised firefly luciferase immediately 5’ of the endogenous stop codon, were created by NEB Hifi assembly. Homology arms of ~1000 bp were included either side of the insert coding sequence. hPSCs were nucleofected with a total of 5ug highly purified plasmid DNA (1:1 molar ratio of linearised HDR template:Cas9 sgRNA) using the 4D-Nucleofector® System and Human Stem Cell Nucleofector Kit 1 (Lonza, VPH-5012), according to the manufacturer’s instructions. 10ml of StemFlex medium was prepared with 1X RevitaCell. Nucleofor reagent was mixed with supplement, then plasmid DNA and left at room temperature. One well of hPSCs at 85% confluence was washed with 1ml DPBS$^{-/-}$ and lifted with 500ul Accutase at 37°C for 4-6 min. The Accutase was neutralized with 1ml RevitaCell™ medium and the culture dissociated into a single cell suspension using a P1000 Gilson. The suspension was transferred to a 15ml falcon and a 10ul aliquot was used for automated counting (Countess FL). $1x10^6$ live cells were transferred to a new 15ml falcon and spun at 100 x *g* for 3 min. The supernatant was aspirated and the cells re-suspended in Nucleofor mix with DNA. Cells were transferred to the cuvette using a P200 Gilson and then transferred to the Nucleofector device for nucleofection (programme B-016). 500ul RevitaCell medium was added to the cuvette using the supplied Pasteur pipette and all contents were gently transferred back to one well of Geltrex-coated 6-well plate containing 2ml RevitaCell™ medium and the plate incubated at 32°C 5% $CO_2$[10]. After 24h the incubator was gradually increased to 37°C, and the media replaced. After 48 h the well was passaged 1 in 3 without RevitaCell and blasticidin was added at selection concentration (5ug/ml) after cells had adhered. Media was replaced every 2 d with media containing 2.5ug/ml blasticidin. After 7 d, sizeable colonies were marked for picking using a P200 Gilson. Each clone was transferred to a single well of a Geltrex-coated 24-well plate for expansion and validation. Cell-permeant TAT-CRE Recombinase (Sigma-Aldrich, SCR508) was used for subsequent removal of the blasticidin cassette, according to the manufacturer's instructions.

*Validation of clock reporter clones*

Clonal lines were expanded until 85% confluent and 300uMfirefly luciferin (Biosynth, #L-8220) was added to the medium. Bioluminescence was checked by taking one 25-min exposure in an ALLIGATOR (Cairn Research) at 37°C and 5% $CO_2$. Positive clones were further validated by PCR (**Supplementary Fig.1a**) and sequencing. The majority of experiments were conducted with biallelic (homozygous) knockin clones for optimal brightness.

*iNeuron differentiation*

NGN2-targeted hPSCs[1,5] were dissociated into single cells with Accutase and plated onto Geltrex-coated dishes at a density of 50,000 cells per $cm^2$. Forward programming was initiated 24h after plating via 100% switch to iNeuron Induction media. On D2, the medium was switched to Neurobasal medium. Where tight osmotic control was required, media switches were osmomatched using 5M NaCl or sucrose and checked with a freezing-point osmometer (Gonotec Osmomat 3000). When indicated, replating was performed on D3 post induction; cells were dissociated into single cells using Accutase and replated for the last time in Neurobasal medium. 50% media changes were performed on D4, D7, D10 and then every 2-3 days. Doxycycline was withdrawn from D6-7 post induction.

*Differentiation of human astrocyte progenitors*

*Bmal1*:Luc astrocyte progenitors were maintained in NSCR EF20 medium[6] with 2.5ug.ml blasticidin added. The monolayer cultures were passaged when confluent using Accutase (split ratio 1:2–1:3). For experiments, progenitors were plated onto Matrigel-coated plates for differentiation into astrocytes (removing blasticidin) as described previously[6]. Differentiation was induced with AstroMED CNTF over 14 d.

*iAstrocyte differentiation*

For iAstrocyte induction, a previously published protocol was used with some modifications[3,11]. At a confluency of 70-80%, targeted S9AB-hiPSCs were dissociated with Accutase into single cells and plated onto Geltrex-coated plates at a density of 20,000 cells per $cm^2$. One day post plating, the medium was changed to FBS enriched medium. Following an additional media change with FBS enriched medium on D1, the medium was gradually changed to FGF enriched medium. On D5 and D6, the medium was changed to full FGF medium. On D8 post plating, half media change was performed to iAstrocyte Maturation medium. On D9 post plating, cells were dissociated with Accutase and replated for final plating on PDL/Geltrex-coated plates or coverslips in iAstrocyte Maturation medium supplemented with 5µM ROCK-I or RevitaCell™. Half media changes were performed every other day. Doxycycline was withdrawn at D14 post induction.

*Human iMicroglia*

Microglia were generated from human iPSCs by opti-ox™ cellular reprogramming (ioMicroglia™, bit.bio Ltd, Cambridge UK). Details of the ioMicroglia cryopreserved product can be found at https://www.bit.bio/products/glial-cells/microglia-wild-type-io1021.

*Human cerebral organoids*

The STEMdiff Cerebral Organoid Kit (StemCell Technologies, 08570) was used to generate cerebral organoids, seeding 2000 cells per embryoid body. After 15 DIV, the cerebral organoids were excised from the Matrigel droplets in which they were embedded during the Kit protocol, with a needle and scalpel under a brightfield microscope, and returned to organoid media. From 30 DIV onwards, organoid media was supplemented with Matrigel dissolved at 2% (v/v). Full media replacement took place twice weekly except during bioluminescent recordings which never exceeded 10 days in succession (**Extended Data Fig.1g**).

*Human ALI-COs*

55-65 DIV cerebral organoids (CO) were cut to 300um slices for culture at the air-liquid interface (ALI), as previously described[12], and collected onto cell culture inserts (Millipore, PICMORG50) with IDM+A beneath. For samples destined for recording on custom-designed MEAs: after slicing, ALI-COs were collected onto cell culture inserts with serum free slice culture medium beneath (SFSCM). After two weeks of ALI-CO culture, the media was switched to BrainPhys Neuronal Medium with N2-A and SM1 (StemCell Technologies, 05793), supplemented with 35ng/ml vitamin C. Media were changed to IDM+A prior to bioluminescence or MEA recordings. All ALI-CO media were supplemented with Pen/Strep (LMB media kitchen, 1:100) and additional 1:1000 (v/v) fungizone (Merck, A2942) or Fungin™ (InvivoGen), and media was refreshed daily until the date of experimentation.

*iAstrocyte validation*

At 14 d post induction, iAstrocytes were incubated for 24h with or without 10 ng/mL IL-1β and TNF-α (PeproTech) in fresh Maturation medium. After stimulation, cells were fixed for immunocytochemistry or lysed for RNA for reverse transcription quantitative PCR (RTqPCR). Total RNA was extracted from iPSCs, and from iAstrocytes at day 7, 14 and 21 post induction using GenElute mammalian total RNA miniprep kit (Sigma-Aldrich) according to the manufacturer's protocol. cDNA synthesis was performed with the Maxima First Strand cDNA Synthesis Kit (Thermo Fisher Scientific). Applied Biosystems SYBR Green PCR Master Mix was used for qPCR. Samples were run on the QuantStudio 6 Flex Real-Time PCR System machine. 3 biological replicates from each sample were analysed in technical duplicates and normalised to the house-keeping gene Porphobilinogen Deaminase 1 (PBGD1). Results were analysed with the ΔΔCt method. See Table 1 for a full list of primer sequences. For RNA-sequencing, total RNA quality and quantity were evaluated using Qubit 4 fluorometer (Invitrogen) with the Qubit RNA BR assay kit (Invitrogen). RNA sequencing libraries were constructed by Novogene UK. Read count abundances were generated from FASTQ files using Kallisto (v0.46.0) by pseudo-alignment to GRCh38.p12, adjusting for GC content bias and bootstrapping 50x. Transcript isoform-level counts were imported to R (v4.2.1), summarized to gene level counts using tximport (v1.26.0), and converted to a DESeq2 object (v1.38) object for normalization. Protein coding genes (total = 18310) were isolated based on GENCODE biotype, and genes with zero counts across all samples were removed (total =1396). Normalized counts were converted to regularized log counts using the rlog function in DESeq2 and principal component analysis (**Supplementary Fig.1b**) was performed using the base R stats package. Heatmaps were generated with ComplexHeatmaps (v2.13.4).

*Primary rodent cortical astrocytes*

Primary mixed glia cultures were used for functional iAstrocyte validation experiments (**Extended Data Fig.2**) and isolated from P0 to P2 neonatal Sprague-Dawley rat forebrains (Charles River Laboratories) as previously described[13-14]. Pups were euthanized according to

“Schedule 1” regulations from the Home Office Animal Procedures Committee UK. Mixed glia cells were maintained for 10 d in culture after which flasks were shaken for 1 h at 260 rpm on an orbital shaker to remove the loosely attached microglia, and then overnight at 260 rpm to dislodge oligodendrocyte precursors. Astrocyte cultures were maintained in glial culture medium (high-glucose DMEM (Sigma-Aldrich) supplemented with 10% FBS, glutamine (Sigma-Aldrich) and 1% Penicillin/Streptomycin) for at least 2 weeks before passaging and plating for MEA recording.

*Bioluminescence recordings*

For bioluminescence experiments, all cultures were retained in their standard 6-, 12- or 24-well culture dishes with (ALI-COs) or without inserts. During their final replating, astrocytes and microglia were plated at confluence to enable contact inhibition and exclude the effects of cell proliferation[15]. 1% P/S was included during all recordings. Doxycycline was included only during recordings that charted bioluminescence during forced differentiation (**Fig.1c**). Anti-fungal reagent was included as a standard for organoid and ALI-CO recordings only. Blasticidin was excluded from all recordings with the exception of early bioluminescence validation of edited clones. In all cases, the final media replacement prior to recording included 300-500uM D-luciferin potassium salt (#L1520 8220, Biosynth), the concentration of which was empirically determined based on optimal bioluminescence in the absence of toxicity. Synchronization with glucocorticoids, media replacement, and any other timing cues were performed as described in figure legends. With the exception of (**Supplementary Fig.3b** and **Supplementary Fig.6**), entrainment with 12h/12 temperature cycles (at temperatures indicated in figures) took place within the ALLIGATOR to enable recording of bioluminescence under cycling conditions, without any disruption to cultures at the transition from cycling to constant conditions. Otherwise, temperature cycling took place in a standard

5% $CO_2$ humidified incubator. For temperature cycling, devices were controlled using a linux-operated temperature program via a laptop connected to the core incubator via an RS232 port. Incubator displays were checked twice daily for acquisition of programmed values, with the temperature log checked at the end of each cycling period to confirm consistent profiles across days. Since standard incubators and the ALLIGATOR system are capable of active heating, but not active cooling, the passive cooling phase always took longer (2-3h) compared to the heating phase (30 min). This profile mimics that which would be observed for diurnal temperature rhythms in humans[16]. All bioluminescence recordings were performed in a light-tight, humidified ALLIGATOR system (Cairn Research)[17] at 5% $CO_2$ in the dark, with 25-min exposures, capturing every 30 min, with EM gain adjusted according to the brightness of the culture.

*Bioluminescence data processing*

ALLIGATOR images were processed in Fiji/ImageJ v2.0-2.14.0/1.54f[18] by first denoising and removing cosmic rays. An average intensity image was created from the image stack in order to create ROIs around single wells, organoids, or ALI-COs. A background ROI was included in the template which was then used to extract luminescence values from the image series. In Microsoft Excel, background subtraction was performed, then a 24-hour moving average was subtracted prior to plotting bioluminescence and detrended bioluminescence data, respectively (**Fig.1b**). Data plotting and rhythm analysis were performed in GraphPad Prism (up to v10.1.1). Non-detrended bioluminescence curves of individual wells, organoids, or ALI-COs were inspected to check for any outliers or exclude non-viable (non-bioluminescent) cultures.

*Drug treatments*

For circadian experiments, all treatments were applied at the constant relevant temperature using a bespoke in-house-engineered temperature-programmable heated platform with heated lid and built-in UPS ('Mr Waffle', courtesy of Martin Kyte, LMB electronics workshop). Mr Waffle was used to transport culture plates to the ALLIGATOR system or MEA platforms to ensure constant temperature conditions were maintained. In all cases, the vehicle control condition included 0.1% DMSO, added at the same volume per well per condition. This concentration has no impact on circadian rhythms in human neural cultures. Where double treatments were applied at 30-min intervals for antagonist studies, working concentrations were doubled such that the final concentration and volume of DMSO was added per well. Compounds used in this study, together with their final media concentrations are listed in the table below:

| **Compound** | **Source** | **Concentration** | **Activity** |
|---|---|---|---|
| Dexamethasone | Sigma | 100nM | Synthetic GR agonist |
| Hydrocortisone | Sigma H0888 | 100nM | Synthetic mimic of cortisol (GR and MR agonist) |
| Corticosterone | Sigma | 100nM | GR and MR agonist |
| Exicorilant (CORT125281)[19] | Corcept Therapeutics | 10uM | Selective GR antagonist |
| Spironolactone | Sigma S3378 | 10uM | Nonspecific MR antagonist |
| Bicuculline | Tocris Bioscience | 40uM | Competitive antagonist at $GABA_A$ receptors |
| CNQX | Tocris Bioscience | 40uM | AMPA and kainate receptor antagonist |

| | | | |
|---|---|---|---|
| GABA | Sigma-Aldrich | 100uM | Primary inhibitory neurotransmitter in brain where concentration is ~1mM |
| Glutamate | Sigma-Aldrich | 100uM | Supratoxic concentration of primary excitatory neurotransmitter in brain |
| LiCl | Sigma-Aldrich | 10-30mM | Clinical overdose level; expected circadian period lengthening |
| Picrotoxin | Sigma-Aldrich | 300uM | Non-competitive antagonist at $GABA_A$ receptors |
| Tetrodotoxin | Tocris Bioscience | 1uM | Potent voltage-gated sodium channel blocker |
| Sucrose | Sigma | +100mOsm | Hyperosmotic stimulus |

*Multiomics timecourse with cerebral organoids*

The paradigm is summarised in **Fig.4a**. With the exception of the bioluminescence and MEA readouts, all organoids had been cultured in suspension (without Matrigel embedding) to exclude the effects of Matrigel on omics data. An extra IHC plate with Matrigel embedding was included as a control for validating cerebral organoid identity; this was harvested at the last time point. $PER2^{BK}$+/+ organoids (N=250) were preconditioned to the experimental temperature (38.5°C) one week before the experiment and maintained at this temperature throughout. At 62 DIV, all organoids underwent 100% media replacement with 100nM hydrocortisone added for robust synchronization. Luciferin (300uM) was included only for the N=6 organoids undergoing bioluminescence, and both this, and the MEA plate (N=6 organoids) were immediately transferred isothermally to their respective recording devices (sampling every 30 min for bioluminescence, and every hour for electrophysiology). For other readouts, plates for each time point were split between two identical incubators according to readout, in order to minimise disruption of the incubator environment with transient door opening. Incubator doors were opened for as short a time as possible and closed gently to avoid vibration. Sampling commenced 24h later to avoid any transient effects of the last resetting cue[8]. For protein, standard RNA, and IHC sampling, plates at each time point were placed immediately on ice. For protein samples, 2 wells (A and B) of a 6-well plate each contained 3 organoids. All media was removed and 1ml ice cold isosmotic DPBS$^{-/-}$ with protease and phosphatase inhibitors (PhosSTOP, Roche, #04906837001 and cOmplete, Roche #4693116001) was added per well. The contents of each well were transferred to a 1.5ml Eppendorf without tituration. Tubes were centrifuged at 500 x *g* for 5 min at 4°C then the supernatant was removed and samples flash-frozen in $LN_2$ and stored at -70°C until further processing. RNA samples had been plated in triplicate as one per well of a 12-well plate (N=3 per time point). For each well in succession, all media was removed and 500ul ice cold isosmotic DPBS$^{-/-}$ (without inhibitors) added. The contents of each well were transferred to a 1.5ml Eppendorf (without tituration), processed and stored as for protein samples. For IHC, organoids were plated as for RNA; all media was removed, the organoid was washed with 500ul ice cold DPBS$^{-/-}$, then 500ul ice cold 4% paraformaldehyde (PFA) was added. The plate was parafilmed and stored for 6h at 4°C. Organoids were then washed 3 x 10min with ice cold isosmotic DPBS$^{-/-}$ at 4°C, with the third wash left in the well and the plate parafilmed and stored at 4°C. For TT-seq samples, organoids were plated as for standard RNA; the plate was immediately placed on an isothermal (38.5°C) heated platform. 1ul 4-thiouridine (4-SU, Cayman Chemical) was added to each well and the media gently titurated 3 x 250ul before the plate went back into the incubator. After exactly 15 min, the plate was transferred to a fume hood at room temperature and the labelling quenched by adding 600ul TRIzol and titurating 5 x to roughly break up the organoid. All material and TRIzol from each well was transferred into a 1.5ml eppendorf and flash frozen in $LN_2$ before storage at -70°C.

*Organoid protein extraction and quantification*

Protein LoBind® Tubes (Eppendorf) were used for all steps. Samples were thawed on ice and centrifuged at 13,000 rpm for 3 min at 2°C to remove remaining supernatant by Gilson. UTH lysis buffer (6M urea, 2M thiourea, 20mM HEPES, adjusted to pH 8) was brought to room temperature and protease and phosphatase inhibitors (PhosSTOP, Roche, #04906837001 and cOmplete, Roche #4693116001) added at 1X just prior to use. 200ul UTH buffer was added per sample, pipetted to mix and incubated at room temperature for 20 min. Samples were sonicated 3 x 30s on/30s off (Bioruptor Plus, Diagenode) and incubated at room temperature for another 20 min. For determination of protein concentration, Pierce 660nm assay (Protein Assay Kit (#22662, Thermo Scientific) was performed with each sample/standard in triplicate in microplate format according to manufacturer's instructions, with bovine serum albumin (BSA) protein standards diluted in UTH buffer plus inhibitors using 2mg/ml BSA stock made from BSA powder. Samples were then prepared for mass spectrometry, with the same protein concentration in the same volume of buffer. For most samples this was 128ug in 213ul, with a few samples needing to be of lower concentration owing to the smaller size of those organoids. 13ul of each sample from each of the time points and each batch (A and B) were pooled, mixed then divided between two tubes to create the $18^{th}$ sample for each TMT run. One sample of the lowest concentration was omitted from the pool in order to submit sufficient quantity of that sample for mass spectrometry. Samples were flash frozen in $LN_2$

and stored at -70°C. Slightly less protein was ultimately submitted for a few samples derived from the smallest organoids, rather than excluding timepoints or submitting small amounts from all samples.

*Protein digestion*

Samples in UTH buffer were reduced with 5mM DTT at 37°C for 40 min and alkylated with 10mM iodoacetamide (IAA) in the dark at room temperature for 30 min. Excess iodoacetamide was quenched by the addition of 5 mM DTT for 10 min. Samples were diluted to 4M urea and incubated with Lys-C (Promega) for 4h at 25°C. Next, the samples were further diluted to 1.5M urea and were digested with trypsin (Promega) overnight, at 30°C. Digestion was stopped by the addition of formic acid (FA) to a final concentration of 0.5%. Any particulate matter was removed by centrifugation at 16000 x *g* for 8 min. Supernatants were desalted using home-made C18 stage tips (3M Empore) filled with 4mg of Oligo R3 (Thermo Scientific) resin. Stage tips were equilibrated with 80% acetonitrile (MeCN)/0.5 %FA followed by 0.5%FA. Bound peptides were eluted with 30-80% MeCN/0.5% FA and lyophilized.

*Tandem mass tag (TMT) labelling*

Dried peptide mixtures (120ug) from each condition was re-suspended in 80 ul of 200mM Hepes, pH 8.5. 40ul TMTpro 18plex reagent (Thermo Fisher Scientific) reconstituted according to manufacturer's instruction, was added to each sample and incubated at room temperature for an hour. The labelling reaction was then terminated by incubation with 8ul 5% hydroxylamine for 30min. The labelled peptides were pooled into one set of TMT sample which was desalted using the same stage tips method as above.

*Off-line high pH reverse-phase peptides fractionation*

About 120ug of the labelled peptides were separated on an off-line, high pressure liquid chromatography (HPLC). The experiment was carried out using XBridge BEH130 C18, 5µm, 2.1 x 150mm (Waters) column with XBridge BEH C18 5µm Van Guard cartridge, connected to an Ultimate 3000 Nano/Capillary LC System (Dionex). Peptides were separated with a gradient of 1-90% B (A: 5% MeCN/10mM ammonium bicarbonate, pH8; B: MeCN/10mM ammonium bicarbonate, pH8, [9:1]) in 60 min at a flow rate of 200µl/min. A total of 54 fractions were collected then combined into 18 fractions and lyophilized. Dried peptides were resuspended in 1% MeCN/0.5% FA and desalted using C18 stage tips and ready for mass spectrometry analysis.

*Enrichment of phosphopeptides and high pH reverse-phase fractionation*

Phosphopeptides were enriched using $TiO_2$ titansphere-chromatography (GL Science Inc. Japan). The rest of the lyophilized peptides were resuspended in a solution of 2M lactic acid in 50% MeCN (loading buffer) and incubated at room temperature for 30 min with $TiO_2$ beads (1:5, peptides: $TiO_2$ beads, w/w), that were prewashed with the loading buffer. Next, the $TiO_2$ beads were centrifuged at 1000 x *g* for 2 min, and the supernatant was transferred into fresh tube that contained $TiO_2$ beads for a second round of enrichment. After incubation, $TiO_2$ beads were loaded onto a C8 stage tips and washed sequentially twice with loading buffer and once with 50% MeCN, 0.1% FA. The bound phosphopeptides were eluted twice with 80ul 0.4M ammonia solution and 50 ul 50% MeCN, 0.1% FA. Samples were then acidified and partially dried down using a SpeedVac (Savant), desalted with a C18 Stage tips and lyophilized. Dried phosphopeptide was fractionated using off-line high pH reverse-phase peptides fractionation as above. The collected fractions were combined into 14 fractions and lyophilized. Dried phosphopeptides were resuspended in 30ul 20% MeCN/0.1% FA; MeCN was removed by vacuum centrifugation, ready for mass spectrometry analysis.

*Mass spectrometry analysis*

The fractionated peptides were analysed by LC-MS/MS using a fully automated Ultimate 3000 RSLC nano System, fitted with a PepMap100 C18 5µm 0.3 x 5 mm nano trap column (Thermo Fisher Scientific) and an Aurora Ultimate TS 75µm x 25cm x 1.7µm C18 column (IonOpticks). Peptides were separated using a binary gradient consisting of buffer A (0.1% FA) and buffer B (80% MeCN, 0.1% FA) at 300 nl/min, column temperature of 40ºC. Eluted peptides were introduced directly via a nanoFlex ion source into an Orbitrap Eclipse mass spectrometer (Thermo Fisher Scientific). The mass spectrometer was operated in real-time database search (RTS) with synchronous-precursor selection (SPS)-MS3 analysis for reporter ion quantification. MS1 spectra were acquired using the following settings: Resolution = 120K; mass range = 400-1400m/z; AGC target = 4e5; MaxIT = 50ms and dynamic exclusion was set at 60s. MS2 analyses were carried out with HCD activation, ion trap detection, AGC = 1e4; MaxIT = 50ms; NCE = 33% and isolation window = 0.7m/z. RTS of MS2 spectrum was set up to search uniprot human reviewed proteome (Jun 2021), with fixed modifications of cysteine carbamidomethylation and TMTpro 16plex at N-terminal and Lys residue. Met-oxidation was set as variable modification. Missed cleavage = 1 and maximum variable modifications = 2. MS3 scans were performed with the close-out function enabled and max number of peptides per protein = 5. The selected precursors were fragmented by HCD and analyzed using the orbitrap with these settings: Isolation window = 0.7 m/z; NCE = 55, orbitrap resolution = 120K; scan range = 110-500 m/z; MaxIT = 300ms and AGC = 1e5.

For phosphoproteomics, multistage activation (MSA) SPS MS3 method was performed on Orbitrap Eclipse. MS1 scans used the same parameters as above except MaxIT was 75ms. MS2 analyses were carried out with CID MSA activation on neutral loss mass 97.9673; detection in the Orbitrap; orbitrap resolution = 60K; AGC = 1e5; MaxIT = 118ms; NCE = 35% and isolation window = 0.7m/z. RTS of MS2 spectrum was set up as for proteomics, with inclusion of phosphor STY as variable modification and maximum variables = 3. In MS3 scans, the selected precursors were fragmented by HCD and analyzed using the orbitrap with these settings: Isolation window = 0.7 m/z; NCE = 55, orbitrap resolution = 120K; scan range = 110-500 m/z; MaxIT = 400ms and AGC = 2e5.

*Raw MS data processing*

The acquired LC-MS/MS raw files, were processed using MaxQuant (Cox and Mann) with the integrated Andromeda search engine (v.2.4.2.0). MS/MS spectra were quantified with reporter ion MS3 for both phosphoproteomics and proteomics from TMTpro 18plex experiments, and searched against Human reviewed UniProt Fasta database (downloaded Jun 2021). Carbamidomethylation of cysteines was set as fixed modification, while methionine oxidation, N-terminal acetylation (protein), and STY phosphorylation (for phosphoproteomics) were set as variable modifications.  Protein quantification requirements were set at 1 unique and razor peptide. Other parameters in MaxQuant were kept as default values.

*Perseus*

MaxQuant output files, proteinGroups.txt and Phospho (STY) sites.txt were then processed with Perseus software (v. 2.0.10.0). After uploading the matrix, the data were filtered to remove identifications from reverse database, identifications with modified peptide only (for protein groups), and common contaminants. The localization probability of phospho (STY) .txt was also filtered to greater or equal to 0.75. Both sets of data with the reporter intensities of '0' were converted to NAN and exported as text file for further data analysis.

*Proteomic and phosphoproteomic analysis*

All subsequent proteomics data processing and analysis was performed in R (v4.3.2) with R Studio (Version 2023.06.2+561) and Cytoscape (v3.10.1). MaxQuant .txt output files were used as a starting point and peptides were filtered to leave only those that were detected in all samples. An initial correction was performed to account for any variable quantities of protein submitted as a result of variation in organoid size. Standard sample loading normalisation was then performed, applying a scaling factor to equalise average intensity across TMT channels, followed by internal reference scaling between runs A and B using the pooled samples. Phosphopeptide data were normalized to their cognate protein intensity before performing rhythmicity analysis. Rhythmicity Analysis Incorporating Non-parametric Methods (RAIN)[20]was used to test for rhythms (TMT runs A and B combined) with period length of 24 h. Whilst each TMT sample from each run included protein from N=3 organoids, the TMT runs were set as biological replicates (N=2). RAIN automatically provides p values corrected for multiple testing using the adaptive Benjamini-Hochberg method; $p<0.05$ was taken as rhythmicity threshold. Oscillation peak phases were taken from RAIN outputs, and represented circadian time relative to the start of sampling, where time 0 is equivalent to GCT 0 (24h post-synchronization with hydrocortisone, 10h prior to the peak of PER2 in parallel bioluminescence recording). Relative amplitude was estimated for each significantly rhythmic protein, where relative amplitude = range/mean abundance. To calculate range, a 24-hour moving average was applied to detrend data before subtracting minimum from maximum values. An arbitrary threshold of 10% is typically used in proteomics analysis to assign biological significance to relative amplitude; in this analysis such a threshold was meaningless, since all rhythmic proteins exceeded it. For gene ontology functional enrichment analysis, GOrilla tool was used[21], comparing target protein list with all detected proteins as background, and setting FDR q-value cutoff at 0.05. REVIGO (0.4) was used to remove redundant terms.

*Multi Electrode Array (MEA) recordings*

For iAstrocyte validation (**Extended Data Fig.2f**), iAstrocytes were tested for their ability enhance electrical activity in iNeurons, and their performance was compared to rat cortical astrocytes. At D3 post induction, iNeurons were dissociated into single cells and replated on PDL/Geltrex coated wells of CytoView MEA 48 (Axion Biosystems) together with D9 iAstrocytes or with rat cortical astrocytes, at a ratio of 1:1. iNeuron monocultures and iAstrocyte monocultures were plated as controls. Co- and mono-cultures were then maintained in 1:1 Neurobasal medium:iAstrocyte Maturation medium for the duration of recordings. Recordings were performed for 20 min at 37°C, 5% $CO_2$ using the Maestro Pro MEA system (Axion Biosystems) at different time points. Circadian recordings from D29 and D64 (**Fig.5**) were performed using the Maestro Pro set to 37°C, 5% $CO_2$ under humidified conditions and sampling for 5 min every hour for at least 96h using the spontaneous Broadband +/- viability mode. Terminal recordings at D74 were conducted as described in **Supplementary Fig.5** using spontaneous Broadband mode. Circadian recordings of D170 whole organoids were performed after re-embedding organoids on Matrigel in a CytoView MEA 6 (Axion Biosystems, N=1 organoid per well). The Maestro Pro was set to 37°C, 5% $CO_2$ under humidified conditions with sampling for 5 min every hour for 96h using the spontaneous Broadband mode. A custom-built device was used for longitudinal MEA recordings of ALI-COs. At 203 DIV, the top surfaces of cell culture inserts were gently flooded with ALI-CO media so that ALI-COs floated. ALI-COs could thus be collected on a spatula and transferred onto custom MEAs (**Fig.5e**) positioned on a new cell culture insert. From this point onwards, media was changed to IDM+A.

*MEA data processing*

Maestro Pro raw data were batch-processed in AxIS Navigator, bandpass filtered 200-5000Hz for high frequency potentials and separately filtered for low frequency potentials from 1-50Hz (where relevant). Key Advanced metrics outputs included number of spikes, bursts, and network bursts in each 5-min sampling, as well as weighted mean firing rate (Hz) and burst duration (s). Spike detection was based on a threshold of +/- 6 SD (in both directions, regardless of polarity). Bursts were defined as a minimum of 5 spikes with a maximum interspike interval of 100ms. Network bursts were defined as a minimum of 50 spikes in a burst with maximum interspike interval of 100ms and minimum 35% of electrodes participating in the burst. For custom-designed MEA analysis, neural data were read into an array containing all the data from all the channels for each .rhs file. This was then bandpass filtered with a Butterworth filter of order 2 between 200 and 5000 Hz, for all the channels sampled at 30kS/s. From these data, peak identification was performed using thresholding in the negative and positive direction. The threshold was set using amplitude cut-off at 6x SD of the channel data. With this, the algorithm was able to count the number of spikes individually. Since all electrodes/channels for a given MEA records field potentials from N=1 ALI-CO, spikes were summed across each channel for each ALI-CO and for each time point, adding the positive and negative spikes together. MEA data for all neural cultures were detrended (subtraction of 24-h moving average) prior to cosinor analysis in GraphPad Prism. The first 24h under constant conditions were excluded from rhythm analyses and extraction of period length.

*Immunocytochemistry (ICC)*

Cells were fixed in 4% PFA for 15 min at room temperature and subsequently washed three times with PBS. The cells were then permeabilized with 0.1% Triton-X-100 (Sigma-Aldrich) for 15 min at room temperature. Then, cells were blocked with 10% goat serum (Abcam) and 0.3% Triton X-100 for 30-45 min at room temperature. Subsequently, cells were incubated with appropriately diluted primary antibodies (GFAP, GeneTex; S100β, GeneTex; Vimentin, Millipore) in 2% goat serum and 0.1% Triton X-100 at 4°C overnight. After three washes with PBS, the cells were incubated for 1 h at room temperature with corresponding goat fluorophore-conjugated secondary antibodies (Alexa Fluor 488, 555 and/or 647; Invitrogen) in PBS supplemented with 1% goat serum. Nuclei were visualized with 4',6-diamidino-2-phenylindole (DAPI; Thermo Fisher Scientific). Cells were mounted on glass slides using Prolong mounting fluid (Invitrogen), dried overnight, and imaged using a Zeiss LSM 710 confocal microscope (Leica).

*Immunohistochemistry of organoid slices*

IHC of organoids (**Supplementary Fig.1c**) was performed as described previously[22].

*Statistical analysis*

Statistical tests were performed using GraphPad Prism (v10.0.3) and R v4.3.2, and are indicated in figure legends. P values are either reported in figures directly, or annotated with asterisks: * $p \leq 0.05$; ** $p \leq 0.01$, *** $p \leq 0.001$; **** $p \leq 0.0001$, ns/NS not significant, $p>0.05$. Unless otherwise stated, smoothed and 24-hour detrended grouped bioluminescence data are presented as solid curves +/- SEM of unsmoothed data as dotted curves (smoothing 6 points either side, 2nd order). Other data are presented as the mean or median ± standard deviation (SD) or 95% confidence interval as stated. For monolayer cultures, N=number of independently edited clones, n=number of wells of cells (technical replicates). Otherwise, N= number of individual organoids or individual organoids from which ALI-COs were generated (biological replicates) and n=number of individual ALI-COs (technical replicates). For comparisons between two groups, the unpaired student t-test was used. For multiple comparisons, one-way analysis of variance (ANOVA) with Dunnett's or Tukey's multiple comparisons test, or Kruskal-Wallis test with Dunn's correction for multiple comparisons were applied. For longitudinal bioluminescence and MEA data, circadian rhythmic parameters were extracted and defined as shown in **Fig.1b**, for each well, organoid, or tissue slice, with tests for circadian rhythmicity performed using cosinor analysis in Prism. Rhythmicity was confirmed if a cosinor fit was significantly preferred over a straight line. The following equation was used as a starting point for curve fitting:

$$y = (mx + c) + ae^{kx} \cos \frac{2\pi x - r}{p}$$

Where m is the baseline, c is the offset from 0 in y-axis, a is the amplitude, k is the damping rate, r is the phase (in radians), and p is the period, which was fixed at an initial value of 24h and constrained within the range 18-36h. The window of analysis for curve fitting excluded the first 24h under constant conditions and circadian phase was then calculated from the timing of peaks of the fitted curve (defined as > 10% above baseline) using Prism's area under the curve analysis of the cosinor fit. The time in hours of the first peak (defined as occurring at >0 h but less than the period value) was converted into circadian phase (dividing it by the period length and multiplying by 24). Conversion to glucocorticoid time (GCT) was performed by subtracting the resultant phase value either from 9.7 (for PER2::LUC data) or from 18.9 (for *Bmal1*:Luc data)[23]. For circular plots of circadian phase in GCT, the Circular package in R[24] was used and circular mean phase is plotted +/- circular SD shown as the width of the wedge. Relative amplitude (RA) was determined from non-detrended data as (peak-trough)/mean baseline, where the difference between the peak and trough during the first 24h under constant conditions was divided by the mean baseline across the first 96 h under constant conditions. Robustness was taken as RA/|k|, where k is the damping rate.

**Neural culture media recipes and supplements**

*iNeurons*

Induction media: DMEM/F12 supplemented with 1% Glutamax, 1% NEAA, 1% N2, 1 % Pen/Strep (1% = 0.5ml), doxycycline (Dox, Sigma-Aldrich, 1-2ug/ml depending on cell density)

Neurobasal medium: Neurobasal supplemented with 1% Glutamax, 2% B-27, 10 ng/mL BDNF (PeproTech), 10 ng/mL NT3 (R&D Systems), 1% Penicillin/Streptomycin, and 1-2 µg/mL Dox.

*Astrocytes*

NSCR EF20 medium: AdDMEM/F12 with 1% N2 supplement, 0.1% B-27, 1% NEAA, 1% Pen/Strep, 1% GlutaMAX (all from Invitrogen), 20 ng/mL FGF-2 (PeproTech), and 20 ng/mL EGF (R&D Systems).

AstroMED-CNTF medium: Neurobasal with 0.2% B-27, 1% NEAA, 1% Pen/Strep, 1% GlutaMAX (all from Invitrogen), and 10 ng/mL CNTF (R&D Systems).

*iAstrocytes*

FBS enriched medium: DMEM/F-12 (Thermo Fisher Scientific), 10% FBS (Sigma-Aldrich) or Hyclone FetalClone III, 1% N2 supplement (Thermo Fisher Scientific), 1% Glutamax

(Thermo Fisher Scientific), 1% Penicillin/Streptomycin (Thermo Fisher Scientific), 1 µg/mL Dox.

FGF enriched medium: Neurobasal (Thermo Fisher Scientific), 2% B-27 (Thermo Fisher Scientific), 1% NEAA (Thermo Fisher Scientific), 1% Glutamax, 1% FBS, 1% Penicillin/Streptomycin, 8 ng/mL FGF (Qkine LTD), 5 ng/mL CNTF (PeproTech), 10 ng/mL BMP4 (Qkine LTD), 1 µg/mL Dox.

Maturation medium: 1:1 DMEM/F-12 and Neurobasal, 1% N2, 1% sodium pyruvate, 1% Glutamax, 5 ng/mL heparin-binding EGF-like growth factor (OriGene Technologies), 10 ng/mL CNTF, 10 ng/mL BMP4

*Organoids and ALI-COs*

Improved Differentiation Media + A (IDM+A): 500ml comprises 250 ml DMEM/F12, 250 ml Neurobasal, 2.5 ml N2 supplement, 10 ml B27+ vitamin A supplement, 125 ul insulin (10mg/ml stock), 500 ul of 50mM 2-ME solution, 5 ml Glutamax supplement, 2.5 ml MEM-NEAA, 5 ml P/S, 750 mg $NaHCO_3$, 5 ml Vitamin C solution (40mM stock). Final glucose concentration in this medium is 17mM.

SFSCM: Neurobasal (Thermo Fisher Scientific, 21103049), 2% B-27 (Thermo Fisher Scientific, 17504044), 1% Glutamax (Thermo Fisher Scientific, 35050038), 0.5% (w/v) glucose

BrainPhys medium: BrainPhys Medium (Stemcell Technologies) supplemented with 2% Neurocult Sm1 Neuronal supplement (Stemcell Technologies) 1% N2 supplement-A (Stemcell Technologies), 350ul of 50ng/ml ascorbic acid, 1% Pen/Strep

**Data availability**

Following peer review, all omics datasets will be deposited in appropriate repositories to facilitate access. All other data supporting the findings of this study are available from the corresponding author on reasonable request.

**Code availability**

Following peer review, custom scripts for proteomics and phosphoproteomics analysis of cerebral organoids will be made freely available online.

**Methods references**
