## Supplementary figures and images for "Circadian clocks in human cerebral organoids"

### Supplementary Figure 1

#### SUPPLEMENTARY FIGURE 1

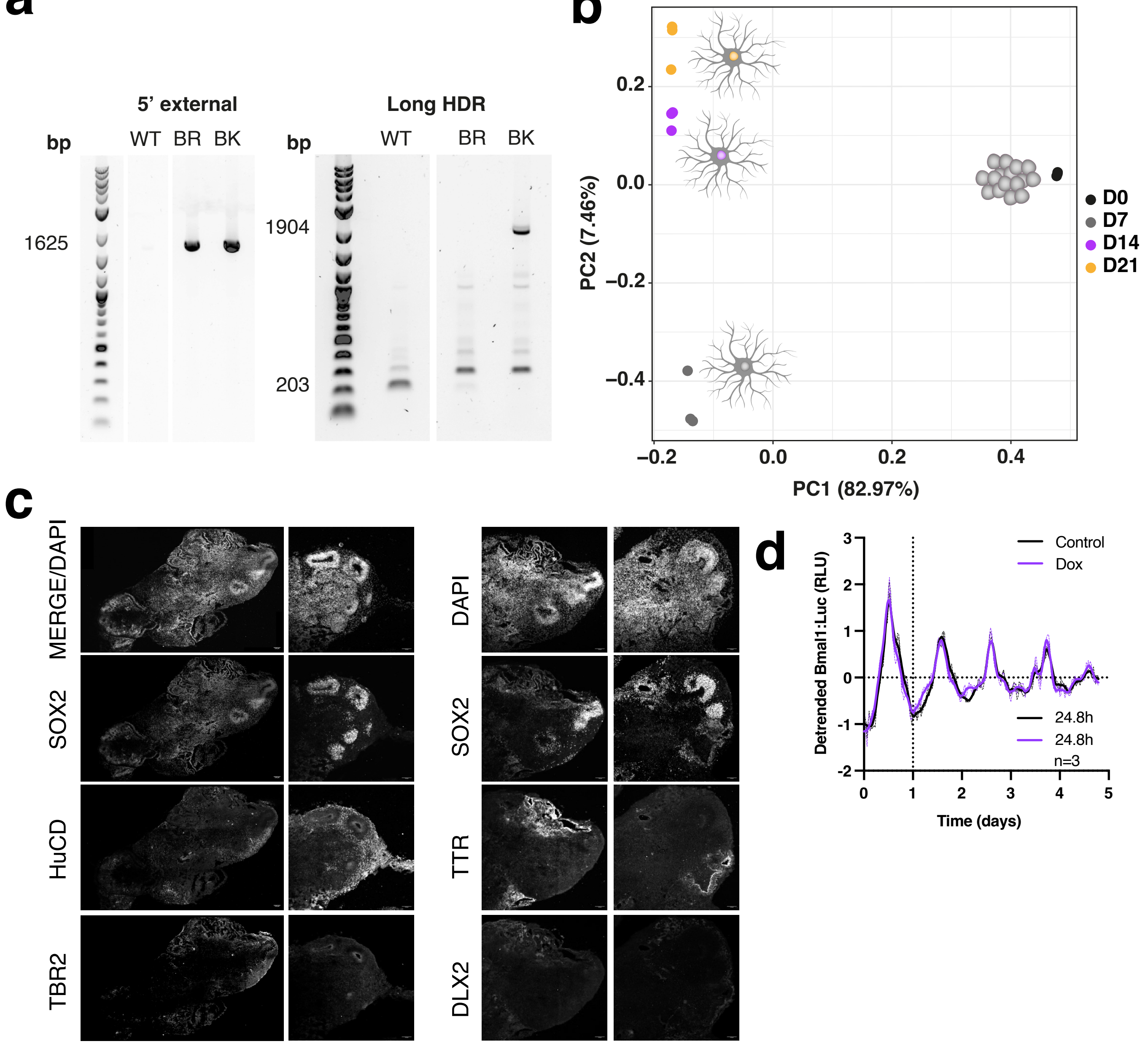

### Supplementary Figure 2

##### SUPPLEMENTARY FIGURE 2

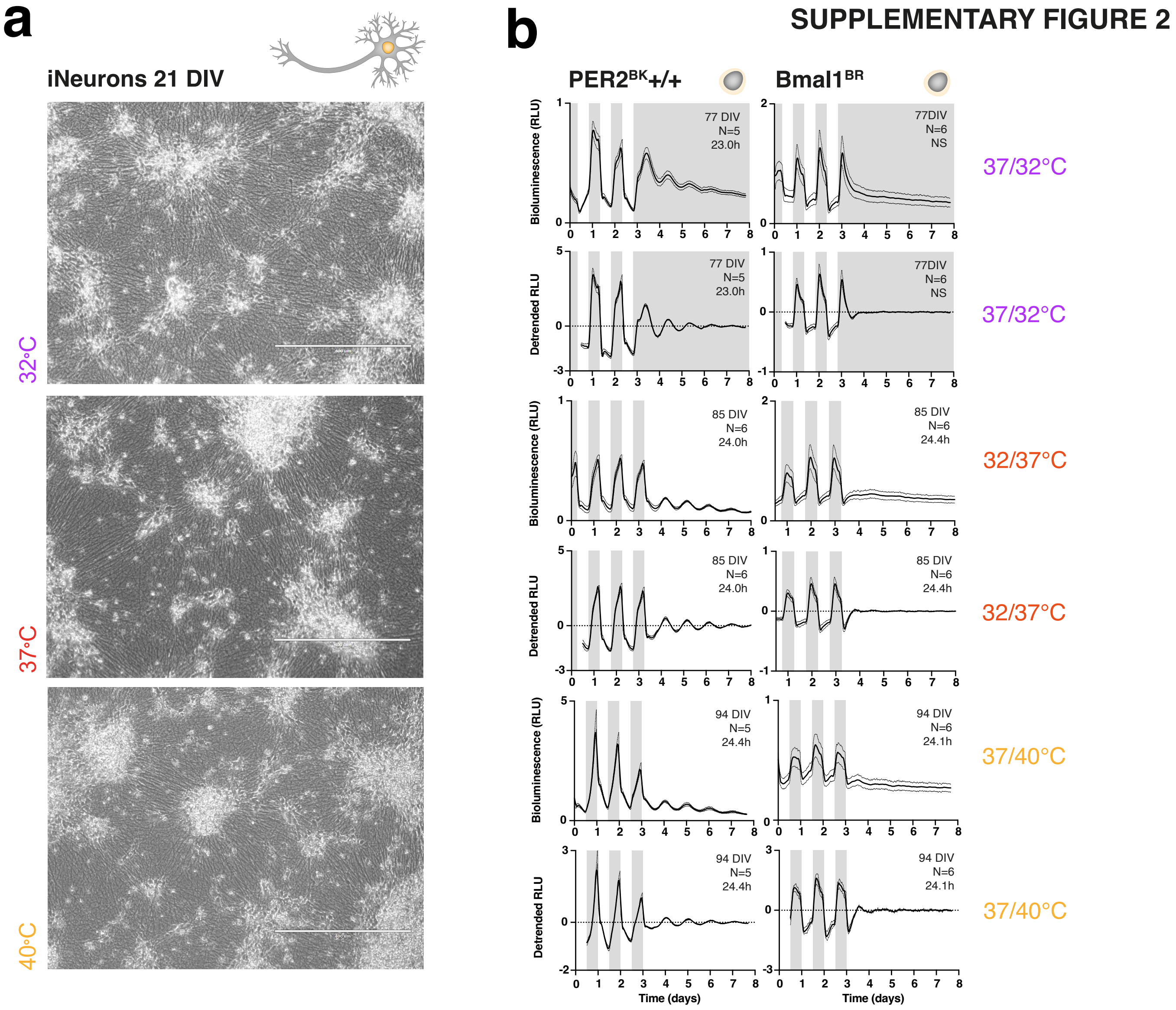

### Supplementary Figure 3

#### SUPPLEMENTARY FIGURE 3

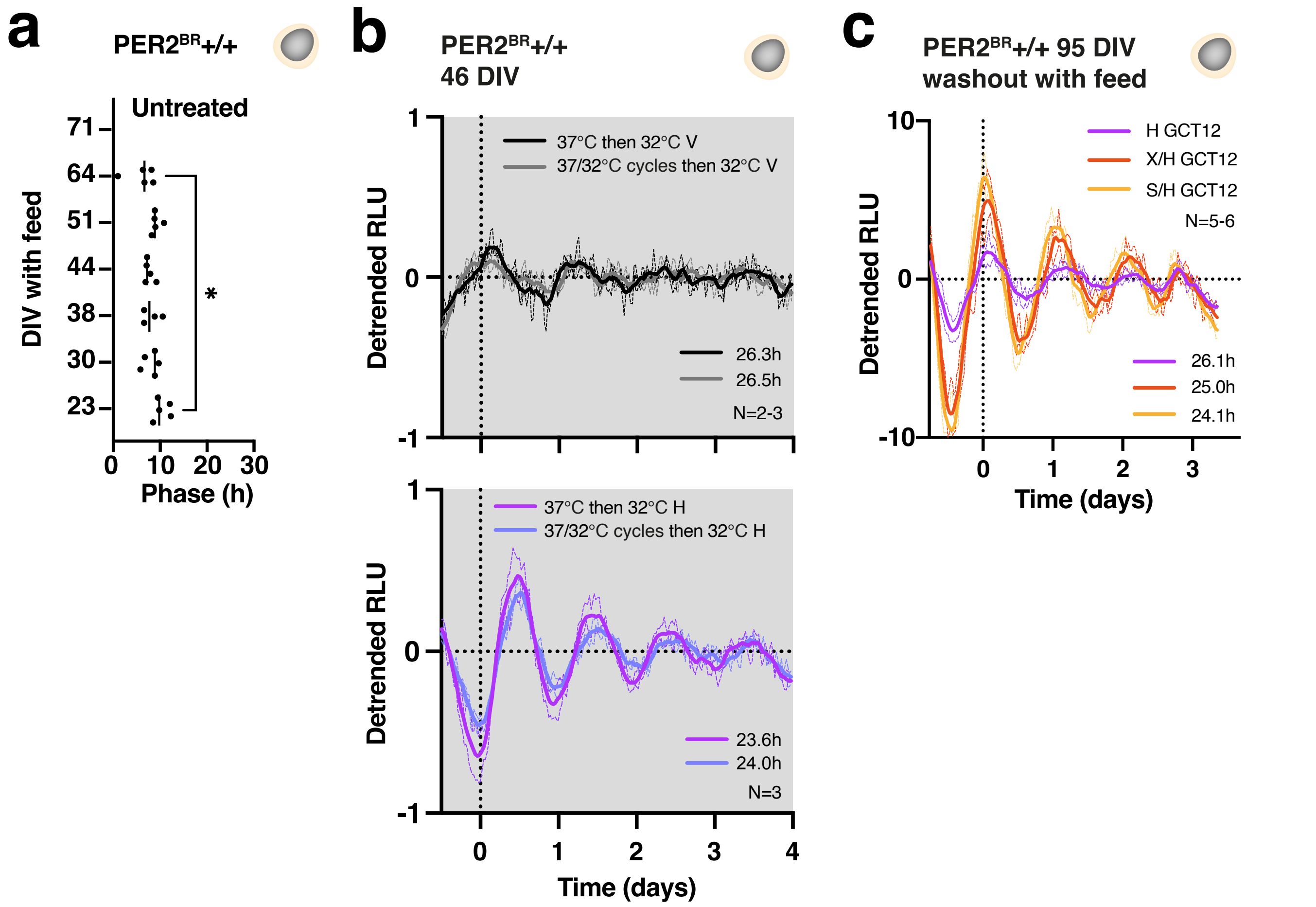

### Supplementary Figure 4

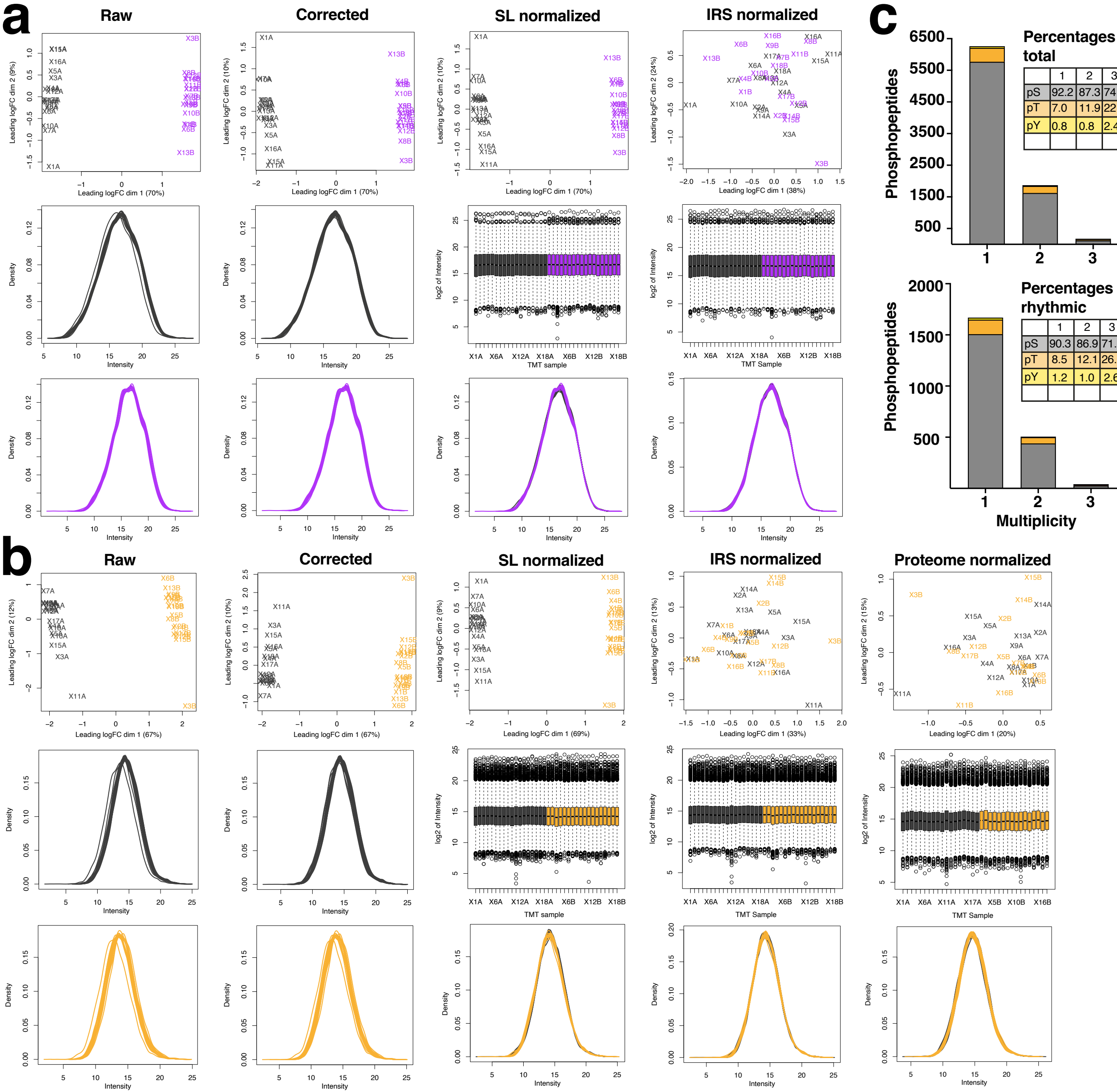


Percentages total

| | 1 | 2 | 3 |
|---|---|---|---|
| pS | 92.2 | 87.3 | 74.7 |
| pT | 7.0 | 11.9 | 22.9 |
| pY | 0.8 | 0.8 | 2.4 |

Percentages rhythmic

| | 1 | 2 | 3 |
|---|---|---|---|
| pS | 90.3 | 86.9 | 71.1 |
| pT | 8.5 | 12.1 | 26.3 |
| pY | 1.2 | 1.0 | 2.6 |

SUPPLEMENTARY FIGURE 4

### Supplementary Figure 5

#### SUPPLEMENTARY FIGURE 5

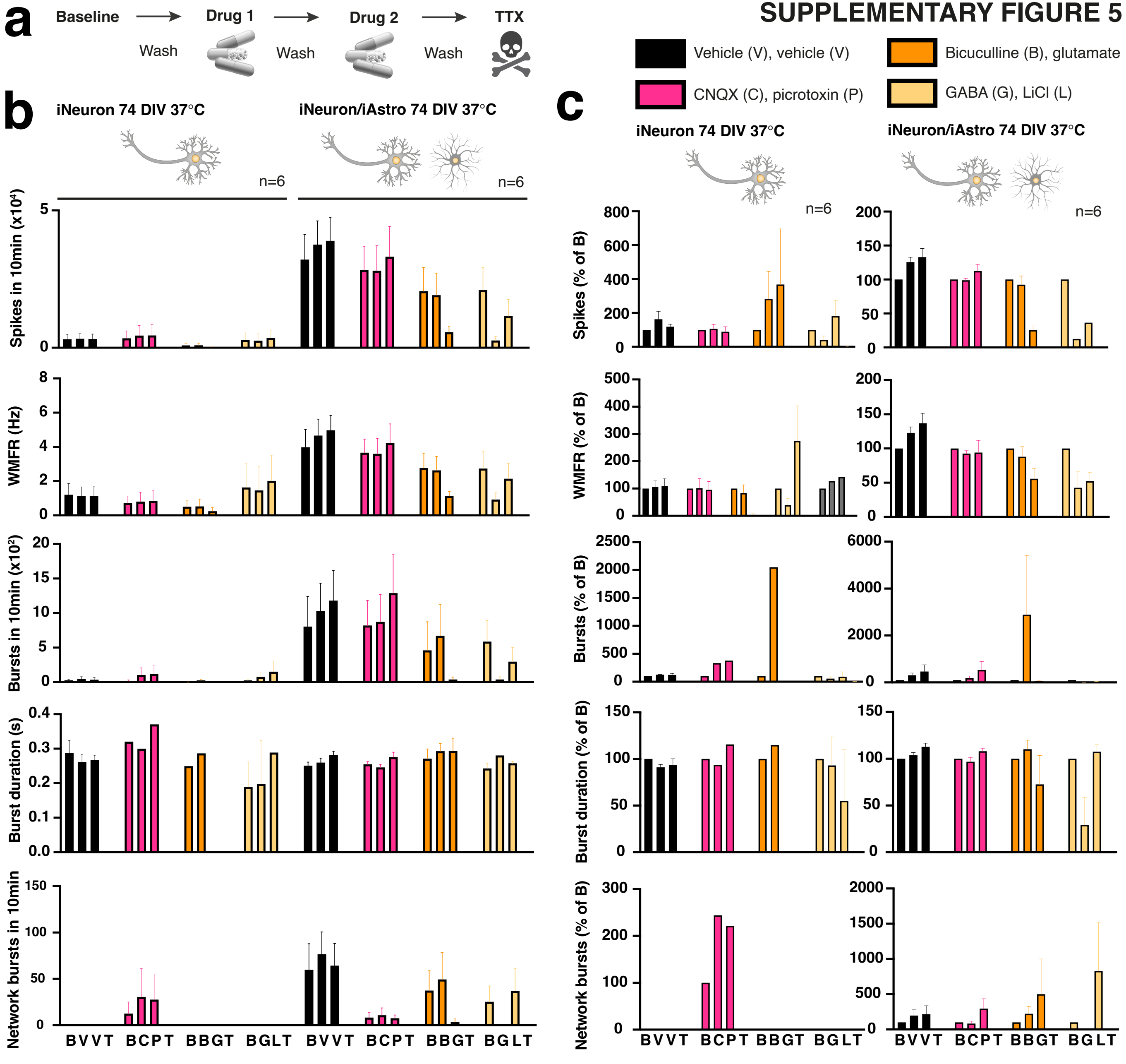

### Supplementary Figure 6

SUPPLEMENTARY FIGURE 6

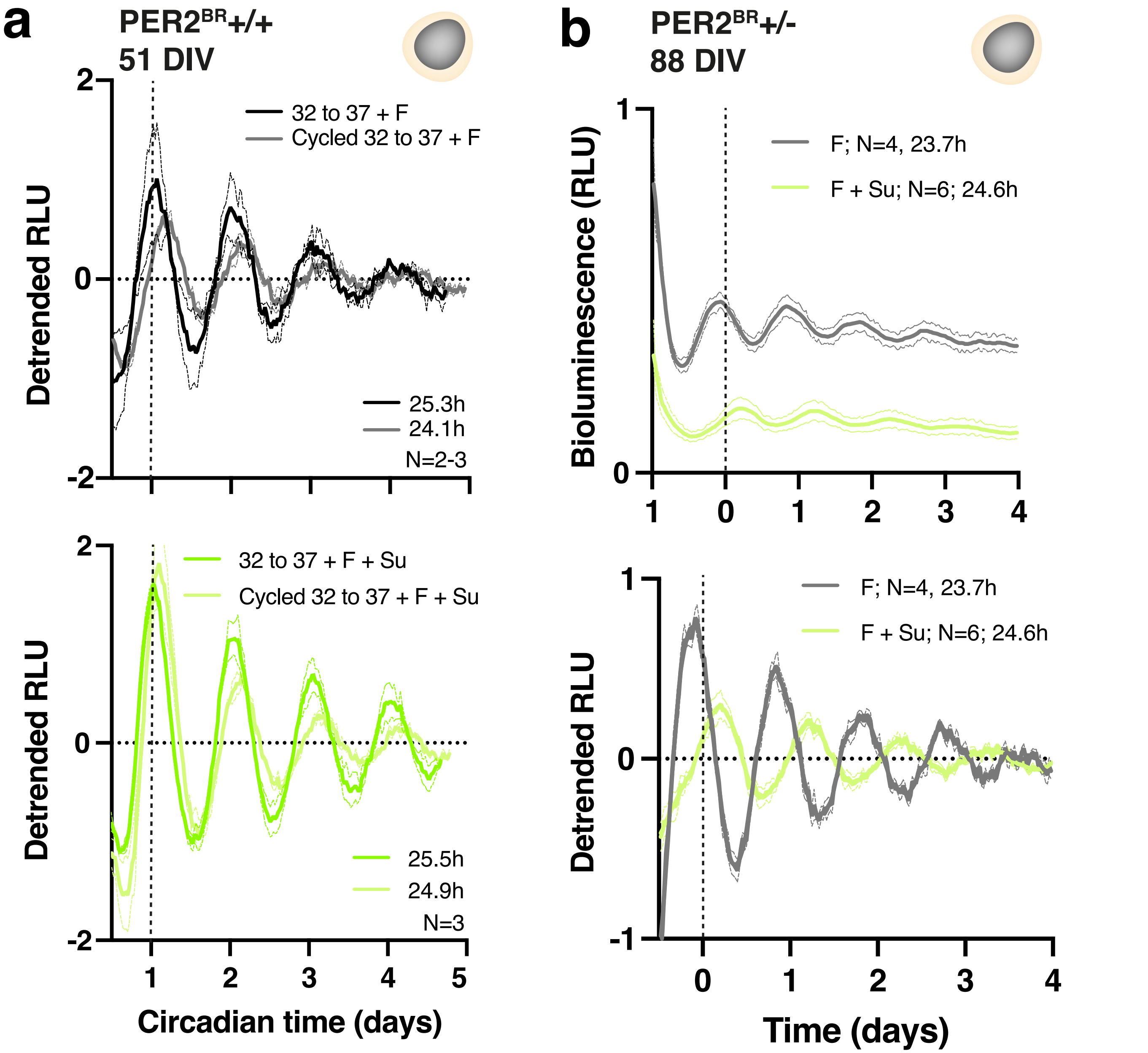
